# An Innovative Mitochondrial-targeted Gene Therapy for Cancer Treatment

**DOI:** 10.1101/2024.03.24.584499

**Authors:** Kai Chen, Patrick Ernst, Seulhee Kim, Yingnan Si, Tanvi Varadkar, Matthew D. Ringel, Xiaoguang “Margaret” Liu, Lufang Zhou

**Affiliations:** Department of Chemical and Biomolecular Engineering, The Ohio State University, Columbus, Ohio, USA; Department of Medicine, University of Alabama at Birmingham, Birmingham, Alabama, USA; Department of Biomedical Engineering, The Ohio State University, Columbus, Ohio, USA; Department of Molecular Medicine and Therapeutics, The Ohio State University, Columbus, Ohio, USA; Comprehensive Cancer Center, The Ohio State University, Columbus, Ohio, USA

## Abstract

Targeting cancer cell mitochondria holds great therapeutic promise, yet current strategies to specifically and effectively destroy cancer mitochondria *in vivo* are limited. Here, we introduce mLumiOpto, an innovative mitochondrial-targeted luminoptogenetics gene therapy designed to directly disrupt the inner mitochondrial membrane (IMM) potential and induce cancer cell death. We synthesize a blue light-gated channelrhodopsin (CoChR) in the IMM and co-express a blue bioluminescence-emitting Nanoluciferase (NLuc) in the cytosol of the same cells. The mLumiOpto genes are selectively delivered to cancer cells *in vivo* by using adeno-associated virus (AAV) carrying a cancer-specific promoter or cancer-targeted monoclonal antibody-tagged exosome-associated AAV. Induction with NLuc luciferin elicits robust endogenous bioluminescence, which activates mitochondrial CoChR, triggering cancer cell IMM permeability disruption, mitochondrial damage, and subsequent cell death. Importantly, mLumiOpto demonstrates remarkable efficacy in reducing tumor burden and killing tumor cells in glioblastoma or triple-negative breast cancer xenografted mouse models. These findings establish mLumiOpto as a novel and promising therapeutic strategy by targeting cancer cell mitochondria *in vivo*.

## Introduction

Mitochondria are the major powerhouses of the cell and vital signaling organelles that regulate key cellular processes essential for maintaining cell growth and function (1). These processes include ATP production (2), redox regulation (3), metabolite generation(^4^), thermogenesis (5), cell division (6), and programmed cell death (7). Importantly, mitochondrial genetics and biochemical metabolisms have been implicated to be associated with various aspects of the cancer cell metastatic cascade, including motility and invasion, modulation of the microenvironment, plasticity, and colonization (8). Given their crucial role in determining cancer cellular function and fate, mitochondria have emerged as a promising target for cancer treatment (9). Over the last decades, numerous mitochondrial-targeted therapies, such as mitocans (10), mitochondriotropics (11), and mitochondriotoxics (12), have been developed to destroy mitochondria and induce cancer cell death. However, these therapies typically target specific signaling pathways or proteins, such as hexokinase (13), Bcl-2 family proteins (14), thiol redox (15), and VDAC/ANT (16), which may undergo unpredictive mutations or develop drug resistance during treatment, thus impairing anti-cancer efficacy (17). More recently, studies reported that butformin (18), a mitochondrial-associated oxidative phosphorylation disruptor, or lonidamine (19), a mitochondrial complex I/II inhibitor, effectively enhanced the anti-tumor efficacy of photodynamic therapy. However, the translation of mitochondrial-targeted therapies to clinics has not yet succeeded.

The mitochondrion is composed of two membranes, a relatively permeable outer membrane and a highly folded and impermeable inner membrane (IMM). Proper mitochondrial function relies on maintaining the electrical potential gradient across the IMM, known as ΔΨ_m_, and a profound and sustained dissipation of ΔΨ_m_ is considered to be a crucial regulatory trigger for cell death (20). Therefore, targeting IMM integrity and disrupting ΔΨ_m_ has generated substantial interest as a potential strategy for cancer treatment. Chemical uncouplers (e.g., FCCP and CCCP) (21,22) or permeability transition pore (mPTP) activators (e.g., Atr and ployP) (23,24) have been used to depolarize ΔΨ_m_. However, because mitochondrial activity is critical for all cells, the lack of specificity of these pharmacological strategies hinders utility *in vivo*. Genetic methods can modulate mitochondrial function in specific tissues but often cause irreversible side effects and do not directly target ΔΨ_m_. Thus, there is currently a dearth of approaches that can directly and dynamically disrupt cancer cell ΔΨ_m_ with greater specificity. Recently, we developed mitochondrial optogenetics (mOpto) by expressing heterologous light-gated Channelrhodopsin 2 (ChR2) in the IMM with a mitochondrial leading sequence (MLS) (25). Importantly, sustained blue light illumination led to irreversible ΔΨ_m_ depolarization and substantial cell death in cells expressing mitochondrial ChR2. However, despite its impressive capability to induce cytotoxicity, mOpto requires external light, which is difficult to penetrate deep tissues in the body, and lacks cancer-specific targeting, limiting its *in vivo* utility and future clinical translation for cancer and other diseases.

To achieve *in vivo* manipulation of mitochondria, we have developed a new-generation optogenetic tool called mitochondrial luminoptogenetics (mLumiOpto, patent US20210205475A1 (26)). This innovative approach harnesses intracellular luminescence as an endogenous light source, eliminating the need for external light stimulation. Specifically, we co-expressed CoChR, a blue light-gated channelrhodopsin from *Chloromonas oogama* (27) in IMM, and an emission spectrum-matched Nanoluciferase (NLuc), a luciferase protein from the deep-sea shrimp *Oplophorus gracilirostris* (28), in the cytosol of the same cells. Additionally, we used a cancer-enhanced promoter (cfos) to maximize selective expression of mLumiOpto genes in tumor cells. Finally, we designed a monoclonal antibody-tagged exosome-associated adeno-associated virus (mAb-Exo-AAV) vehicle to achieve targeted delivery of mLumiOpto genes to tumors *in vivo*. We hypothesized that the mLumiOpto approach enables optogenetics-mediated cancer mitochondrial depolarization and cytotoxicity with the synthesized intracellular bioluminescence (NLuc). Furthermore, we hypothesized that use of the mAb-Exo-AAV will facilitate cancer-specific gene delivery and functional expression of mLumiOpto, allowing for targeted elimination of cancer cells with minimal side effects *via* synergizing mitochondrial depolarization-mediated direct cell death/apoptosis and mAb or AAV-mediated *in situ* immunity within the tumor microenvironment (TME).

To test these hypotheses, we examined the ability of mLumiOpto to induce mitochondrial depolarization and cytotoxicity across different cancer cell types. Then the cancer-specific surface binding, internalization, transduction efficiency, biodistribution, and tumor-specific expression of mLumiOpto were assessed *in vitro* and *in vivo*. The therapeutic efficacy of mLumiOpto, delivered with free AAV and mAb-Exo-AAV, was evaluated in preclinical mouse models carrying GBM and TNBC xenograft, respectively. GBM is a malignant and heterogeneous brain tumor subtype accounting for ∼50% of primary intracranial gliomas, while TNBC represents a highly aggressive breast cancer subtype associated with poor survival. Our results demonstrated that mLumiOpto effectively induces cancer cell death and significantly reduces tumor burden without impairing normal organs or tissues.

## Results

### mLumiOpto development and optimization

We first constructed the mLumiOpto plasmid to co-express light-gated rhodopsin in the IMM and an emission spectrum-matched luciferase in the cytoplasm of the same cancer cells. Specifically, we synthesized the *NLuc-2A-ABCB10-CoChR* expressing plasmids by cloning CoChR (peak λ_ex_ = 470 nm) and NLuc (peak λ_em_ = 460 nm), which were fused through a cleavable 2A linker, into a pcDNA3.0 expression vector (Fig. 1a). We utilized CoChR instead of the more commonly used ChR2 due to its much higher (∼10-fold) photocurrent(27) and efficiency in inducing mOpto-mediated ΔΨ_m_ depolarization (Fig. 1b). NLuc was chosen for mLumiOpto as it generates much brighter bioluminescence compared to other discovered blue light-emitting luciferases such as *Renilla luciferase* (RLuc) (29) (Fig. 1c) and *Gaussia luciferase* (GLuc) (30) (data not shown) when coupled with ViviRen, an engineered luciferin. We then fused the ABCB10 mitochondrial leading sequence (MLS) (31) to the N-terminal of CoChR to facilitate mitochondrial expression. Consistent with our previous studies (25), ABCB10 MLS led to high-level and mitochondrial-specific CoChR expression across various tumor cell lines, including HeLa (Manders overlap coefficient, M1=0.99±0.03) and TNBC MDA-MB-231 (M1=0.98±0.1) (Fig. 1d), as revealed by the strong overlap between eYFP (green, fused with CoChR) and MitoTracker (red, a mitochondrial indicator). Confocal microscopy imaging also confirmed the co-expression of NLuc (fused with eGFP) and CoChR (fused with mCherry) in the transfected MDA-MB-231 cells (Fig. 1e).

**Figure 1.**
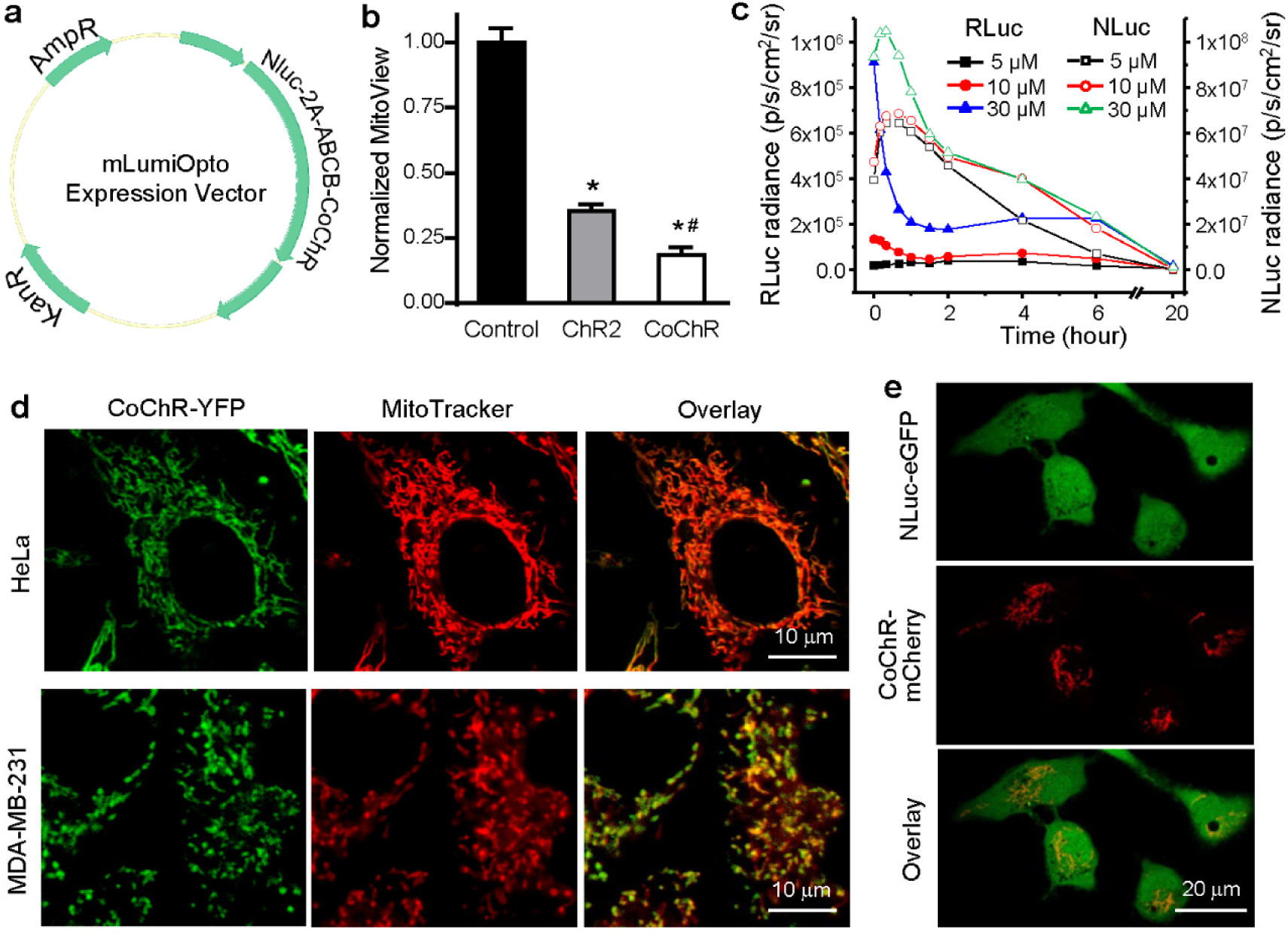
Development of mLumiOpto technology. (**a**) Map of mLumiOpto expression vector carrying the luciferase (i.e., NLuc) and mitochondrial rhodopsin (i.e., ABCB-CoChR) genes linked *via* a cleavable 2A linker. (**b**) Photostimulation with LED light (0.5 mW/mm^2^, 24 hours) caused more severe mitochondrial depolarization (measured by MitoView, a voltage-sensitive fluorescent dye) in cells expressing ABCB-CoChR compared to those expressing ABCB-ChR2. (**c**) IVIS imaging revealed that NLuc emitted much stronger luminescence than RLuc when coupled with luciferin ViviRen. (**d**) ABCB10 mitochondrial leading sequence resulted in high-level, mitochondrial-specific CoChR expression in HeLa and MDA-MB-231 cells, as demonstrated by the strong overlap of eYFP (green, fused to CoChR) and MitoTracker dye (red, a mitochondrial marker). (**e**) Representative confocal images demonstrated the co-expression of NLuc-GFP and CoChR-mCherry in mLumiOpto plasmid-transfected TNBC MDA-MB-231 cells. *: *P*<0.05 vs. control (mock transfected). #: *P*<0.05 vs. ChR2. n=4/group.

### mLumiOpto induces cancer cell mitochondrial depolarization and cytotoxicity

To validate the functionality and efficiency of mLumiOpto to mediate cancer mitochondrial depolarization, MDA-MB-231 TNBC cells were transfected with *NLuc-2A-ABCB10-CoChR* plasmid and then induced with varying doses (0-100 μM) of ViviRen. ViviRen elicited intracellular NLuc luminescent responses (Fig. 2a) and ΔΨ_m_ depolarization (Fig. 2b) in a dose-dependent manner, confirming the functional expression of NLuc and CoChR proteins and the capability of mLumiOpto to depolarize mitochondria. Prolonged exposure to ViviRen (48 hours) caused a dose-dependent reduction in cell viability in mLumiOpto-transfected MDA-MB-231 cells (Fig. 2c). This mLumiOpto-mediated cytotoxicity was further observed in multiple human GBM (U251 and U87) and TNBC (BT-20 and MDA-MB-468) cell lines, exhibiting substantial cell death with induction of ViviRen (Fig. 2d). Importantly, at the tested levels, neither mLumiOpto plasmid transfection nor ViviRen induction alone had a significant effect on cancer cell ΔΨ_m_ (Fig. S1) and cell growth and viability (Fig. 2c-2d).

**Figure 2.**
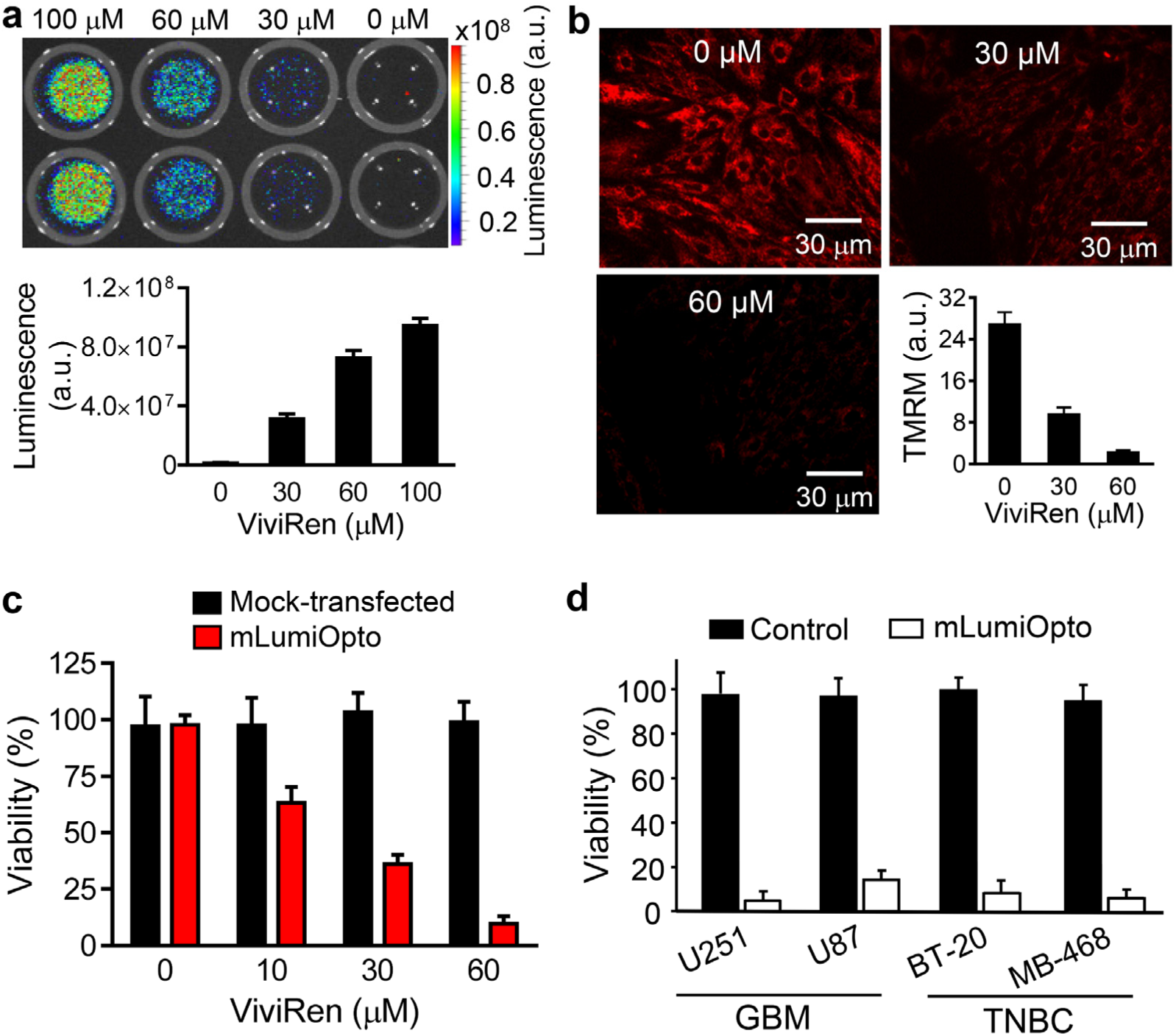
*In vitro* characterization of mLumiOpto technology. (**a**) IVIS imaging showed that ViviRen elicited dose-dependent bioluminescent responses in mLumiOpto plasmid-transfected cancer cells. (**b**) Confocal imaging showed that ViviRen induced mLumOpto-mediated mitochondrial depolarization in a dose-dependent manner, as measured by MitoView 633, a mitochondrial membrane potential-dependent fluorescent dye (red). (**c**) ViviRen caused dose-dependent cell death in mLumiOpto-transfected TNBC (MDA-MB-231) cells but had no obvious cytotoxic effect in mock-transfected cells. (**d**) mLumiOpto caused substantial cytotoxicity in multiple cancer cell lines, including GBM cells (U251 and U87) and TNBC cells (BT-20 and MDA-MB-468). n=4-6/group.

We next investigated the underlying mechanisms of mLumiOpto-mediated cancer cell death. Specifically, we examined the effect of inhibitors for apoptosis (Z-VAD-FMK), necroptosis (Necrostatin-1, Nec-1), or autophagy (Bafilomycin A, BfA) on TNBC MDA-MB-231 cells expressing mLumiOpto genes that were treated with ViviRen for 48 hours. Our results showed that Z-VAD-FMK significantly improved mLumiOpto-mediated cell death, while Nec-1 and BfA had no noticeable effect on cell viability (Fig. 3a). Similar results were observed in GBM U251 cells: while the apoptosis inhibitor effectively attenuated mLumOpto-induced cytotoxicity, the effect of necrosis and autophagy inhibitors were trivial (Fig. S2). These findings suggest that mLumiOpto-induced cell death is mainly due to caspase-dependent apoptosis, rather than necroptosis or autophagy. To further investigate the intrinsic or extrinsic nature of apoptosis, we co-treated mLumiOpto-expressing cells with ViviRen and caspase-specific inhibitors. Both Z-DEVD-FMK and Z-LEHD-FMK significantly attenuated mLumiOpto-induced cytotoxicity, while Z-IETD-FMK had no obvious effect (Fig. 3a), indicating that the activated apoptotic pathway is intrinsic. The apoptotic cell death was further confirmed by a significant increase in caspase-3 activity (Fig. 3b) and TUNEL-positive cells (Fig. 3c) in mLumiOpto-treated cells compared to control cells (i.e., mock-transfected, transfection only, or ViviRen only). We also observed apparent cytochrome C release in mLumiOpto-treated cells, revealed by the punctated/diffused cytochrome C staining disparate from the TOM20 immunostaining (Fig. 3d). Collectively, our results demonstrate that mLumiOpto-induced cytotoxicity is mainly through the mitochondrial-dependent intrinsic apoptosis pathway.

**Figure 3.**
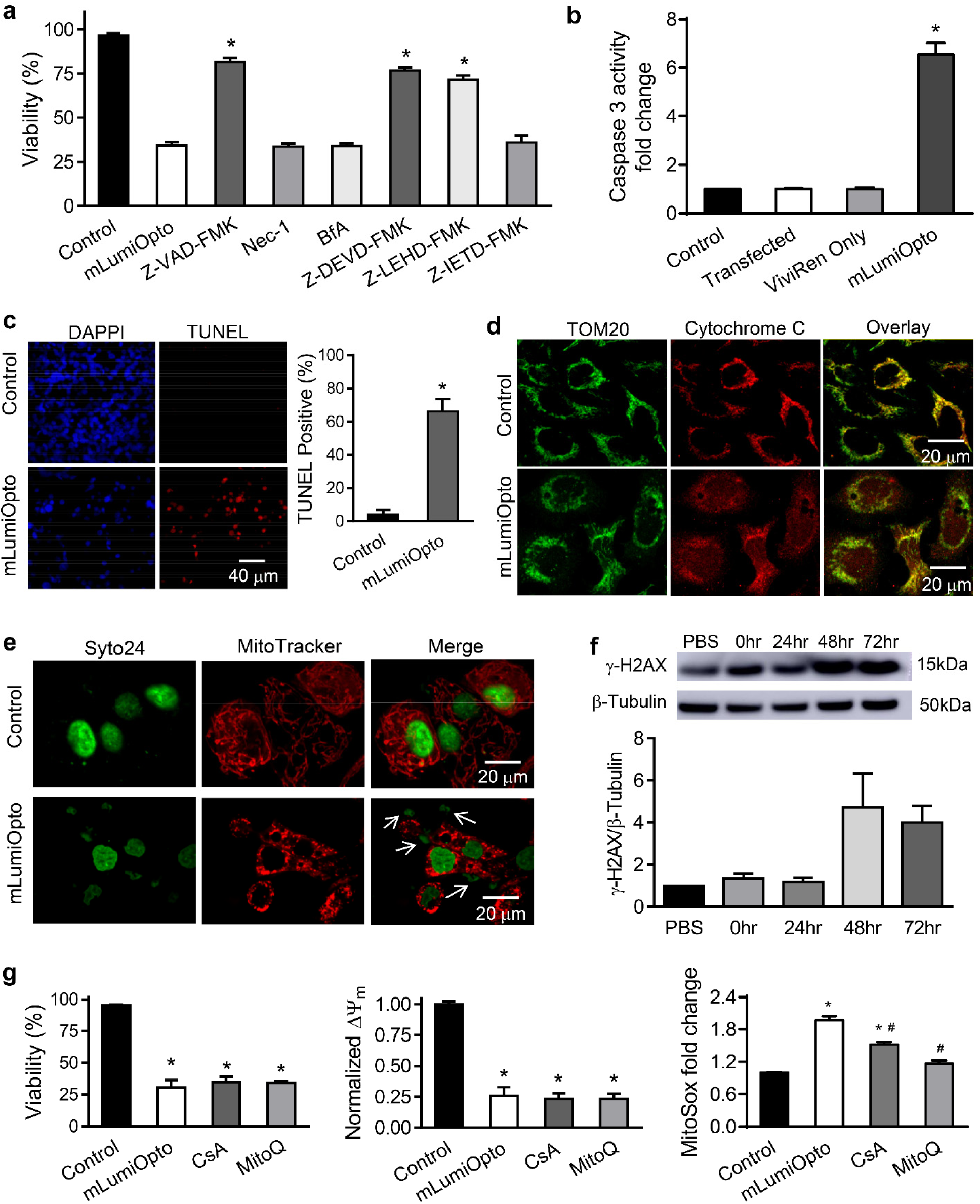
Mechanistic pathways underlying mLumiOpto-mediated cytotoxicity. (**a**) Z-VAD-FMK (a pan-caspase inhibitor), Z-DEVD-FMK (a caspase 3 inhibitor) and Z-LEHD-FMK (a caspase 9 inhibitor) significantly alleviated mLumiOpto-mediated cancer cell (MDA-MB-231) death, whereas Nec-1 (a necrosis inhibitor), BfA (an autophagy inhibitor) and Z-IETD-FMK (a caspase 8 inhibitor) had no effect. (**b**) Caspase 3 activity increased significantly (> 6 folds) in mLumiOpto-treated cells compared to controls (e.g., PBS, ViviRen only, and transfected only). (**c**) The number of TUNEL-positive cells was significantly higher in mLumiOpto-treated cancer cells compared to the control. (**d**) Representative immunostaining showed apparent cytochrome C release in mLumiOpto-treated cancer cells but not in control. (**e**) Syto23 staining revealed substantial DNA damage induced by mLumiOpto. (**f**) Western blotting confirmed DNA damage as indicated by increased expression of ʏ-H2AX in TNBC cells. (**g**) CsA (an mPTP opening inhibitor) and MitoQ (a mitochondrial-specific antioxidant) had no significant effect on mLumiOpto-mediated cancer cell mitochondrial depolarization and cytotoxicity. *: *P*<0.05 vs. mLumiOpto (a) or control (b, c and g). #: *P*<0.05 vs. mLumiOpto. n=4/group.

We conducted further investigations into the effects of mLumiOpto treatment on cancer cells. Our results showed that nuclear condensation and fragmentation occurred in the treated cancer cells, as observed through Syto24 staining (Fig. 3e). Western blot analysis confirmed the occurrence of DNA damage, as evidenced by the increased expression of γ-H2AX, a marker for DNA double-strand breaks, in the cancer cells treated with mLumiOpto for 48-72 hours (Fig. 3f). We also examined the role of mitochondrial permeability transition pore (mPTP) opening and excessive mitochondrial-derived reactive oxygen species (mtROS) in mLumiOpto-mediated cytotoxicity, as these factors are known to contribute to mitochondrial-mediated cell death. While the mPTP inhibitor Cyclosporin A (CsA) and mitochondrial antioxidant MitoQ effectively reduced mtROS levels, neither of these agents significantly improved mLumiOpto-induced ΔΨ_m_ depolarization or cell death (Fig. 3g). Our data indicates that mLumiOpto induces cancer cell cytotoxicity mainly through mitochondrial-mediated intrinsic apoptosis and DNA damage, which are independent of mPTP or mtROS.

### AAV construction and characterization for in vivo gene delivery

To deliver the synthesized mLumOpto genes to cancer cells *in vivo*, we constructed an AAV expression vector using a commercial hybrid serotype AAV-DJ/8 with a heparin-binding domain mutation, which has shown high infection efficiency *in vivo* (32–35). Additionally, we cloned and utilized the cfos promoter (36) to enhance cancer-selective gene expression. AAV was produced in a stirred-tank bioreactor and purified using ion-exchange liquid chromatography as we recently reported (37). The size (∼20 nm) and morphology of purified AAV DJ8 were verified using TEM (Fig. 4a). Western blotting confirmed the expression of three viral capsid proteins, including VP1 (87 kDa), VP2 (73 kDa), and VP3 (62 kDa) (Fig. 4b). Importantly, our data showed that cfos mediated remarkably higher (more than 50 folds) GFP expression in cancer cells compared to non-cancerous H9C2 myoblasts (Fig. S3a), indicating the cancer selectivity of cfos promoter. Moreover, ViviRen triggered robust luminescence (Fig. 4c), along with substantial mitochondrial depolarization (Fig. 4d) in AAV-transduced human GBM U87 cells, demonstrating functional expression of mLumiopto proteins. Consistent with observed mitochondrial collapse, ViviRen induction caused dramatic cell death in various GMB cell lines transduced with mLumiOpto AAV, including the drug-resistant U251-TMZ cells (Fig. 4e). On the contrary, ViviRen (30 μM) had no significant effect on the viability of mLumiOpto AAV co-cultured non-cancerous cells such as HTori-3 thyroid cells (Fig. S3b) and 184B5 mammary epithelial cells (Fig. S3c).

**Figure 4.**
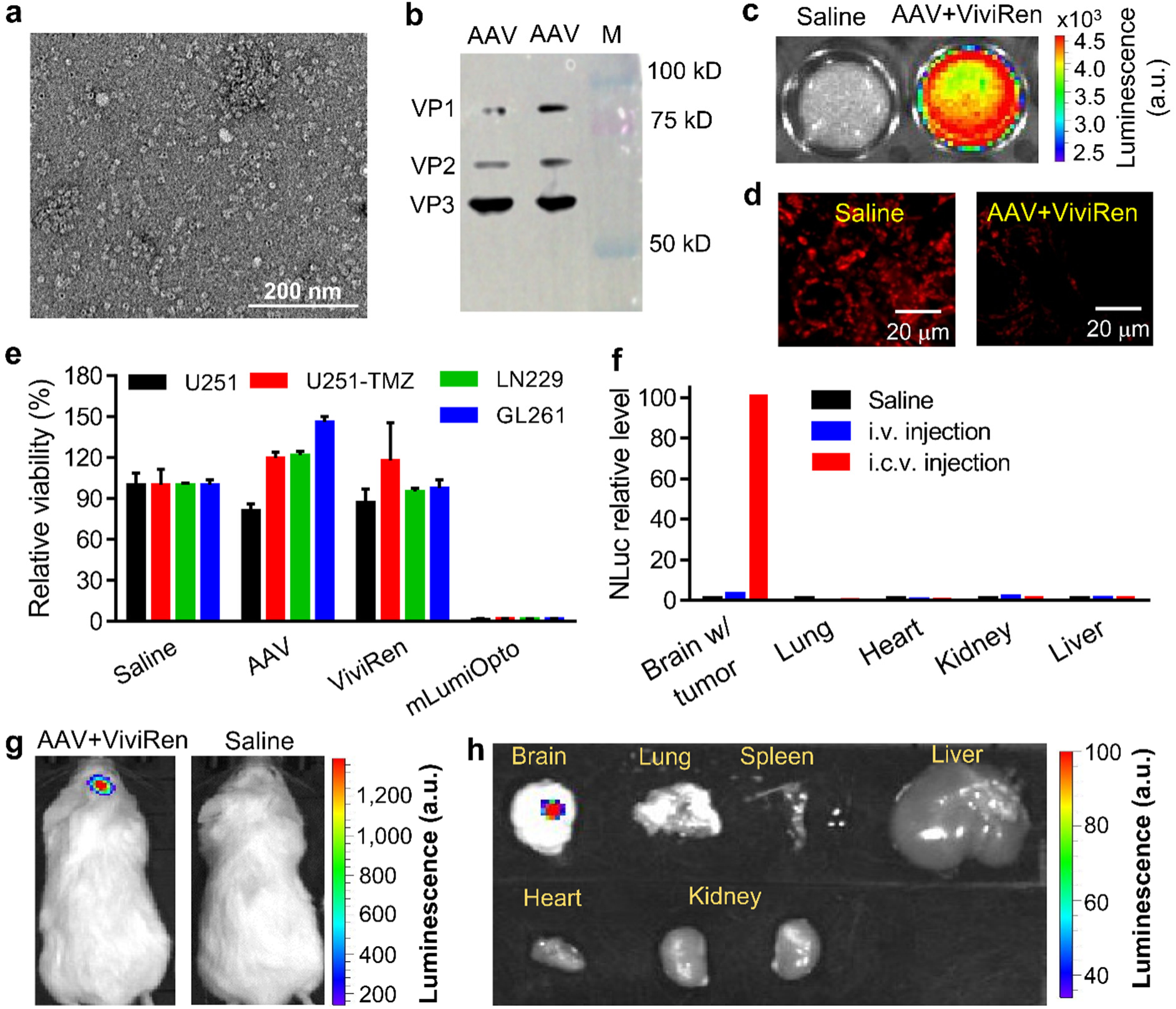
Characterization of AAV-mediated mLumiOpto gene delivery in GBM xenograft mouse model. (**a**) Transmission electron microscopy imaging showed AAV DJ8 particles with the correct morphology and size (∼20 nm). (**b**) Western blotting confirmed the presence of AAV viral capsid proteins VP1 (87 kDa), VP2 (73 kDa), and VP3 (62 kDa). (**c**) ViviRen (30 µM) induced strong NLuc luminescence in mLumiOpto AAV-transduced GBM U87 cells. (**d**) mLumiOpto (AAV+ViviRen) induced severe mitochondrial depolarization in GBM U87 cells, as measured by MitoView fluorescence. (**e**) mLumiOpto killed 90-99% of GBM U251, U251-TMZ, LN229, and GL261 cells within 72 hours, while AAV or ViviRen alone had no significant cytotoxic effect on these cells. (**f**) Intracerebroventricular (i.c.v.) AAV injection led to remarkably higher (∼33-35 folds) mLumiOpto gene (NLuc) expression in GBM tumors compared to intravenous (i.v.) injection. (**g**) ViviRen elicited strong luminescence in mLumiOpto AAV-transduced intracranial GBM xenografts. (**h**) *Ex vivo* IVIS imaging confirmed mLumiOpto expression in GBM xenografts but not in normal organs. n=4-6/group.

### AAV delivered mLumiOpto for GBM treatment

We next assessed the *in vivo* anti-cancer efficacy of mLumiOpto using GBM xenograft mouse model. To test the *in vivo* gene delivery efficiency of AAV, we performed and compared the intravenous (i.v.) and intracerebroventricular (i.c.v.) injections of AAV into GBM xenograft model. qRT-PCR analysis of the harvested tumor and important organs (lung, heart, kidney and liver) revealed that i.c.v. injection achieved GBM tumor-specific mLumiOpto gene delivery and remarkably higher levels of NLuc (Fig. 4f) and CoChR (Fig. S4) expression than i.v. injection. Moreover, live-animal IVIS imaging (Fig. 4g) and *ex vivo* imaging of the isolated organs (Fig. 4h) confirmed the functional expression of mLumiOpto genes in AAV (*via* i.c.v. injection) transduced GBM xenografts. Consequently, direct intracranial administration was utilized for the following anti-GBM efficacy studies.

To evaluate the *in vivo* GBM treatment efficacy of mLumiOpto, the xenografted mice were randomly divided into four groups (n=8-10/group) and i.c.v. administrated with saline (control), AAV only, mLumiOpto 1 (AAV dose: 0.5×10^11^ vg/mouse), and mLumiOpto 2 (AAV dose: 1.0×10^11^ vg/mouse), respectively, on Days 7 and 14. Mice in the mLumiOpto groups received ViviRen through tail vein injection daily for 3 consecutive days following each AAV administration. Our data showed that mLumiOpto significantly prolonged the survival of GBM xenografted mice compared to control groups (Fig. 5a). Body weight profiles were similar across all groups (Fig. 5b). The H&E staining of tumor parafilm section slides showed that mLumiOpto treatment significantly reduced GBM tumor burden (Fig. 5c). IHC staining of tumor slides with antibodies of cleaved caspase 3 (cCasp3) and Ki67 indicated mLumiOpto induced apoptosis and inhibited cell proliferation (Fig. 5d). Moreover, immunofluorescence assay revealed evident cytochrome C release in the treated group, implying mLumiOpto induced GMB mitochondrial depolarization and injury *in vivo* (Fig. S5). No damage in normal organs of the brain, heart, lung, liver, spleen, and kidney was detected (Fig. 5e). IVIS imaging performed on Day 28 (i.e., 10 days after the last ViviRen administration) revealed that the GBM tumor volumes in mLumiOpto groups were reduced by >10 folds compared to control groups (Fig. 5f and Fig. S6). Endpoint MRI imaging on Day 44 (26 days post-treatment) confirmed the reduction of GBM tumor burden in the brain with mLumiOpto treatment compared to controls (Fig. 5g). The ViviRen only had no effect on GBM tumor and mouse body weight and no toxicity to normal organs (data not shown). It is worth noting that the anti-tumor efficacy is comparable between mLumiOpto 1 and mLumiOpto 2 groups (0.5 vs. 1.0×10^11^ vg/mouse), suggesting that AAV dose and administration schedule may need further optimization in the future.

**Figure 5.**
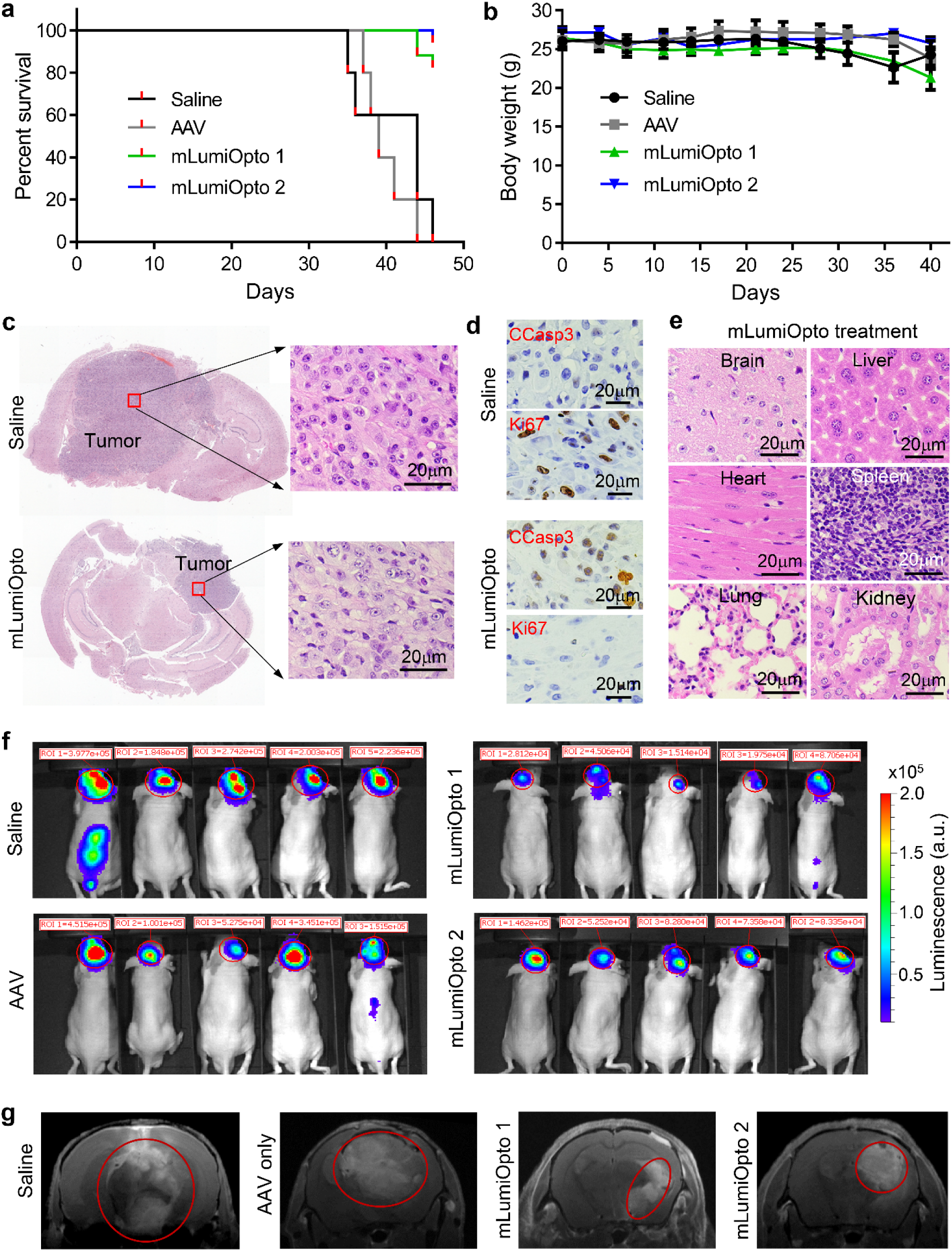
Evaluations of anti-cancer efficacy of AAV-delivered mLumiOpto in GBM xenograft mouse model. (**a**) mLumiOpto treatment significantly extended the survival of GBM xenograft mice. (**b**) Body weight trends were similar among the control and treatment groups. (**c**) H&E staining of the parafilm section slides of GBM xenograft demonstrated tumor burden reduction by mLumiOpto. (**d**) IHC staining of tumor slides with antibodies of cleaved caspase 3 and Ki67 indicated apoptosis-induced cell death and inhibition of proliferation, respectively, post-treatment. (**e**) H&E staining did not detect obvious injury or toxicity in normal organs of the treatment group. (**f**) IVIS imaging revealed >10-fold reduction in GBM tumor size in mLumiOpto treatment groups compared to control (saline and AAV only) groups. (**g**) MRI images taken in the late stage of the survival study (i.e., 44 days post-cell implantation or 23 days after the last ViviRen injection) confirmed a significant reduction of GBM tumor burden in mLumiOpto treatment groups. n=8-10/group.

In addition to assessing the efficacy of mLumiOpto in treating aggressive U87 xenografts, we also investigated its effectiveness in heterogeneous GBM using PDX models. Firstly, subcutaneous GBM PDX xenografts were generated in mice (n=8/group), followed by i.v. injection of AAV and ViviRen. The PDX tumor weight in the treatment group was 40% lower than that in the saline group (Fig. 6a), while body weight profiles remained similar (Fig. 6b). Histological examination of major organs, including the brain, heart, lung, liver, spleen, pancreas, and kidney, did not reveal any signs of inflammation, apoptosis, or necrosis (Fig. 6c), indicating the tumor specificity and safety of AAV-delivered mLumiOpto. Secondly, intracranial GBM PDX xenograft mouse models were established, and mice were treated with i.c.v. injection of AAV and ViviRen. MRI images at the endpoint (14 weeks post-implantation) showed a significant reduction in GBM PDX tumor burden in the mLumiOpto-treated group compared to the saline group (Fig. 6d).

**Figure 6.**
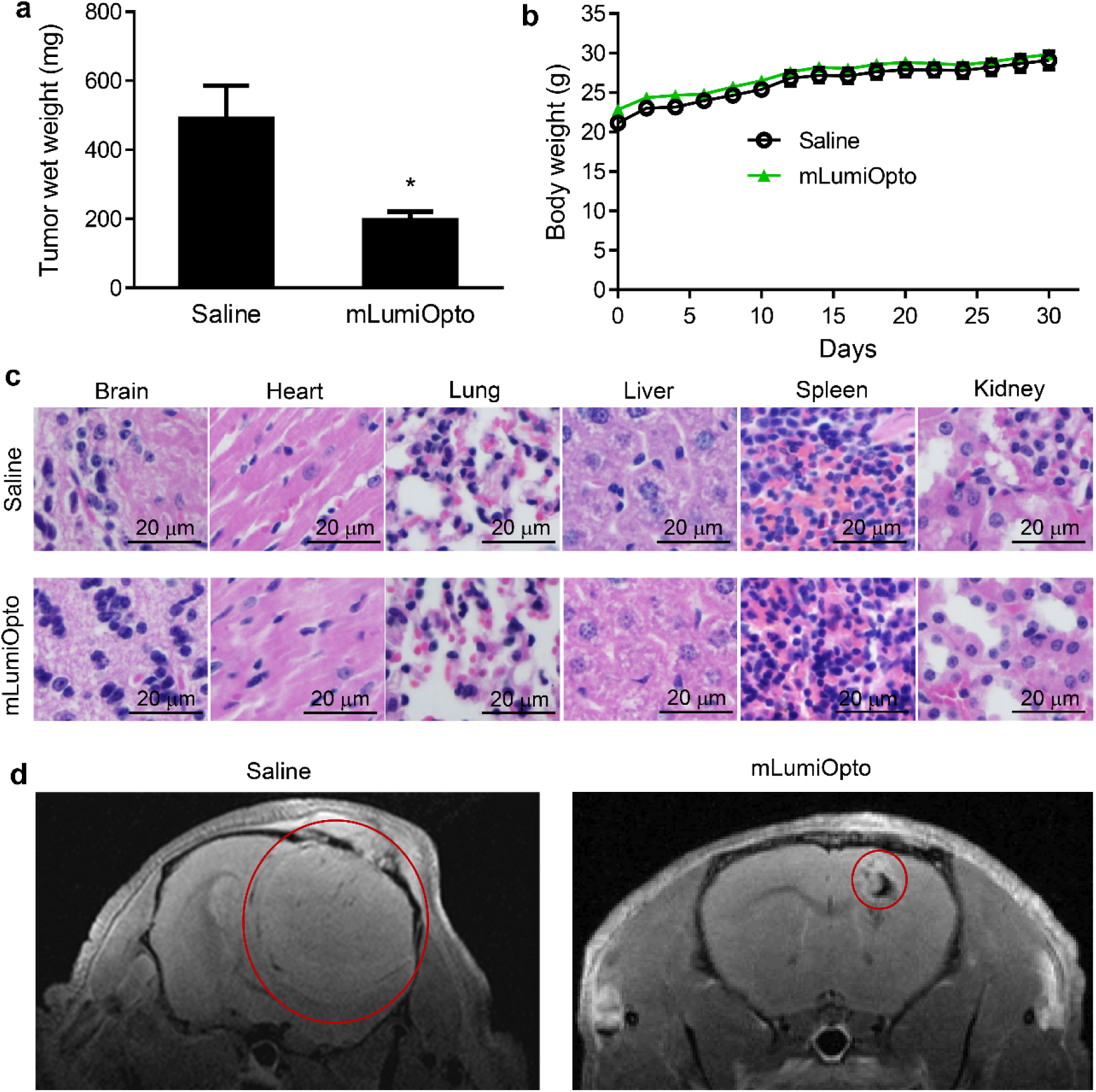
Evaluation of anti-cancer efficacy of mLumiOpto in GBM PDX xenograft mouse model. (**a**) mLumiOpto treatment caused a significant decrease in tumor wet weight compared to the control group. (**b**) Body weight did not exhibit significant differences between groups. (**c**) H&E staining did not detect any damage in normal organs (brain, heart, lung, liver, spleen, pancreas, kidney) of the treatment group. *: P<0.05 vs. Saline. n=5-8/group. (**d**) MRI images of intracranially xenografted GBM PDX at the endpoint (14 weeks post PDX xenograft) showed that GBM tumor burden was significantly reduced by mLumiOpto treatment. n=4-5/group.

### Construction and characterization of mAb-Exo-AAV for targeting gene delivery in vivo

To achieve highly efficient and targeted delivery of mLumiOpto genes to cancer cells *in vivo*, we developed a monoclonal antibody (mAb)-tagged exosome-associated adeno-associated virus (mAb-Exo-AAV) delivery vehicle. Briefly, we first produced high-quality and high-yield Exo-AAVs using viral production cells 2.0 in a 2-L stirred-tank bioreactor. We then surface-tagged the purified Exo-AAV with a model anti-epidermal growth factor receptor (EGFR) mAb, cetuximab, *via* mPEG-DSPE linker as we previously described (38), generating TNBC-targeting mAb-Exo-AAV (Fig. 7a). NanoSight showed the size distribution of mAb-Exo-AAV is at 133.4±74 nm (Fig. 7b), transmission electron microscope (TEM) image confirmed the morphology and particle size of Exo-AAV (Fig. 7c). The packed AAV exhibited high AAV packing rates of 20-50 vg/ptc of exosomes with a mean value of ∼30 vg/ptc. Alexa Fluor 488 dye was used to label mAb and detect the tagging ratio, which revealed that there were 20-100 copies of mAb on each Exo-AAV particle. Flow cytometry assay showed that anti-EGFR mAb-Exo-AAV had strong surface binding to EGFR^+^ TNBC MDA-MB-231 (>60%) and MDA-MB-468 (>95%) cells at room temperature (Fig. 7d). Furthermore, confocal microscope imaging confirmed that Cy5.5-labeled mAb-Exo-AAV bound to the surface of TNBC MDA-MB-468 (Fig. 7e) cells within 20 minutes post incubation and internalized within 2 hours. We also observed the accumulation of Cy5.5-labeled AAV in >95% of cells 30 minutes after transduction (Fig. 7f), suggesting high transduction efficiency. Finally, ViviRen triggered strong NLuc luminescence in mAb-Exo-AAV-transduced MDA-MB-231 cells, demonstrating the functional expression of the NLuc protein in cancer cells (Fig. 7g).

**Figure 7.**
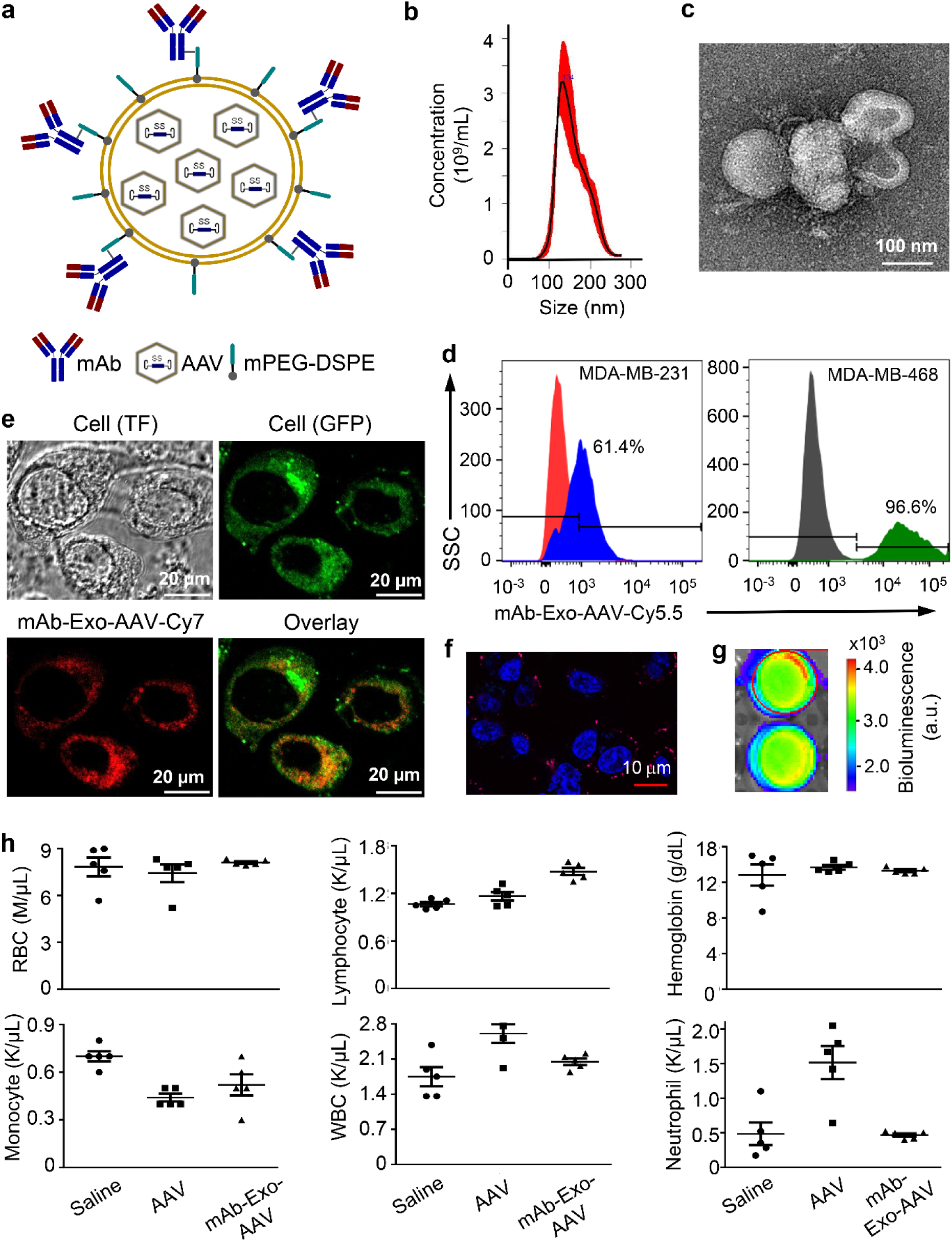
Construction and *in vitro* characterization of mAb-Exo-AAV. (**a**) Schematic description of the mAb-Exo-AAV structure. The anti-EGFR mAb is tagged to the surface of Exo-AAV *via* the DMPE-PEG-NHS linker. (**b**) The size distribution of mAb-Exo-AAV was determined by NanoSight. (**c**) TEM imaging showed the expected morphology and size of mAb-Exo-AAV. (**d**) Western blotting confirmed the expression of exosome biomarkers, including CD81, CD63 and HSP70. (**e**) Flow cytometry showed that mAb-Exo-AAV bound strongly to TNBC cells (MDA-MB-468 and MDA-MB-231). (**f**) Confocal imaging confirmed that mAb-Exo-AAV-Cy5.5 (red) is bound to the surface and internalized by TNBC MDA-MB-468 cells. (**g**) Confocal imaging showed that mAb-Exo-AAV-Cy5.5 (red) transduced over 95% of TNBC cells and started accumulating around the nucleus (blue) 30 minutes after incubation. (**h**) IVIS imaging confirmed that ViviRen induction elicited bright NLuc luminescence in mAb-Exo-AAV-transduced MDA-MB-231 cells. (**i**) Complete blood cell count demonstrated low peripheral toxicity of mAb-Exo-AAV in BALB/cJ mice. n=5/group.

To evaluate the potential immune response of mAb-Exo-AAV in the TME and its effect on general immunity, we administered mAb-Exo-AAV (2×10^13^ ptc/kg-BW), free AAV (2×10^13^ ptc/kg-BW), and saline (control) to healthy wild-type BALB/cJ mice *via* tail vein. Two weeks later, whole blood samples were collected and analyzed using an automatic complete blood count. Our data showed that mAb-Exo-AAV had no significant effect on blood cell counts except for lymphocytes, which were approximately 30% higher than those in the control groups (Fig. 7h) but remained within the normal range (0.9-9.3 K/µL). In contrast, free AAV led to an increase in white blood cells and neutrophils while reducing monocytes (Fig. 7h). These data suggest that mAb-Exo-AAV causes less peripheral immunity compared to free AAV, but this feature requires further investigation.

### mAb-Exo-AAV mLumiOpto inhibits tumor growth in preclinical TNBC models

We next conducted a comprehensive evaluation of mAb-Exo-AAV-delivered mLumiOpto for tumor treatment, including tolerated dosage, tumor targeting, biodistribution, and anti-tumor efficacy. To investigate the tolerated dosage and potential toxicity, various doses of mAb-Exo-AAV were i.v. injected into C57BL/6J wild-type mice, followed by administration of ViviRen three days later to induce intracellular bioluminescence. The normal body weight of the mice indicated no major toxicity of mLumiOpto at the tested dosages (Fig. S7a). Whole blood analysis reported normal cell counts of erythrocytes, leukocytes and thrombocytes (Fig. S7b-d). Furthermore, H&E staining of major organs did not reveal any apparent inflammation, apoptosis or necrosis (Fig. S7e). Consistent with histology analysis, echocardiograph revealed normal cardiac function in the mice (Fig. S7f). No toxicity or damage was observed in the liver and kidney of BALB/cJ mice treated with mAb-Exo-AAV, as indicated by the similar serum levels of alanine transaminase (ALT), aspartate aminotransferase (AST), aspartate aminotransferase (BUN), and creatinine (Fig. S7g). Taken together, our findings suggest that the mAb-Exo-AAV is a safe vehicle for delivering the mLumiOpto gene and that the mLumiOpto technology has minimal toxicity in healthy animals.

To assess tumor-specific targeting and biodistribution, TNBC MDA-MB-231 xenograft NSG female mice were i.v. administrated with anti-EGFR mAb-Exo-AAV. Live-animal IVIS imaging demonstrated strong NLuc luminescence overlaying with the tumor (Fig. 8a), indicating that mAb-Exo-AAV specifically targeted TNBC tumors in mouse models. Consistent with *in vivo* imaging, the *ex vivo* imaging detected bright Cy7 fluorescence only in tumors but not in normal organs such as brain, heart, lung, liver, kidney, and spleen (Fig. 8b). To further examine the biodistribution of mLumiOpto delivered with mAb-Exo-AAV, the MDA-MB-231 xenograft mice were administered with saline (control) or anti-EGFR mAb-Exo-AAV. Three days later, mice were sacrificed to harvest tumors and major organs. Both qRT-PCR analysis (Fig. 8c) and semi-quantitative PCR (Fig. S8) revealed significant CoChR expression in tumor tissue of mAb-Exo-AAV mice but not in that of control mice. Importantly, the expression of CoChR gene in normal organs (e.g., heart, brain, lung, kidney, intestine, spleen, pancreas, stomach, colon, and liver) of the mAb-Exo-AAV mice was undetectable (Fig. 8d). Collectively, these data demonstrated that anti-EGFR mAb-Exo-AAV could specifically bind to tumors and deliver mLumiOpto genes to EGFR^+^ TNBC *in vivo*.

**Figure 8.**
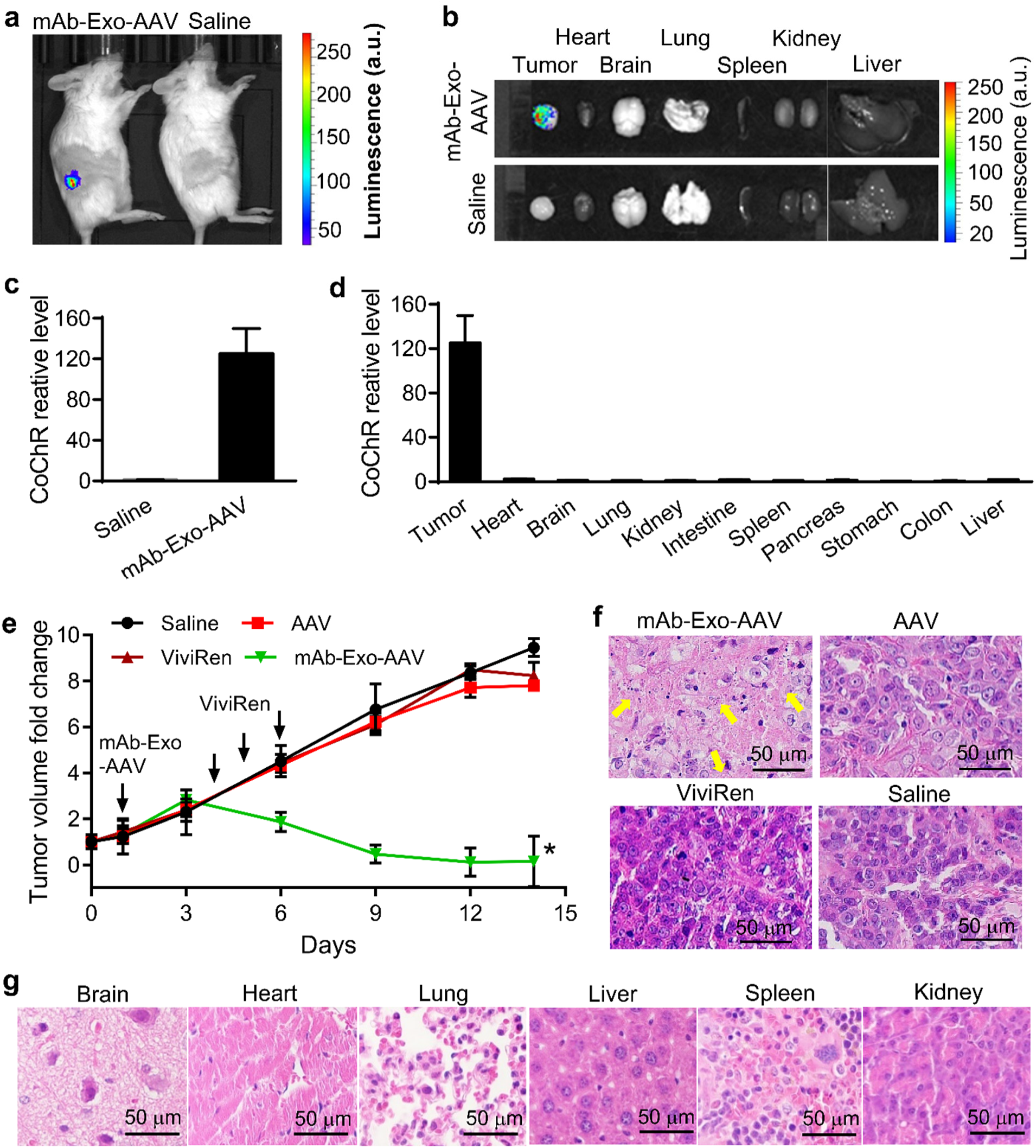
Evaluation of anti-cancer efficacy of mAb-Exo-AAV-delivered mLumiOpto in human TNBC MDA-MB-231 xenograft mouse model. (**a**) Live animal IVIS imaging revealed the overlay of NLuc luminescence (mAb-Exo-AAV) with TNBC xenograft. (**b**) *Ex vivo* IVIS imaging showed the accumulation of mAb-Exo-AAV in TNBC xenografts but not in normal organs (e.g., heart, brain, lung, spleen, kidney and liver). (**c**) CoChR gene expression was high in the tumor of mLumiOpto mAb-Exo-AAV mice but not in that of Saline mice. (**d**) CoChR expression was high in TNBC tumors but undetectable in normal tissues of mAb-Exo-AAV mice. (**e**) The tumor volume was reduced in mLumiOpto-treated human TNBC xenograft mice compared to control groups (saline, AAV and ViviRen alone). (**f**) H&E staining revealed obvious cell death in tumors of mLumiOpto-treated (i.e., mAb-Exo-AAV+ViviRen) mice. (**g**) H&E staining showed no injury in normal organs of mLumiOpto-treated mice. *: *P*<0.05 vs. controls. n=6/group.

To assess the anti-cancer efficacy, we i.v. injected anti-EGFR mAb-Exo-AAV into TNBC MDA-MB-231 xenografts when the tumor volume reached 25-50 mm^3^. The tumor-bearing mice given saline, free AAV, or ViviRen only were used as controls. Remarkably, the TNBC tumor stopped growing shortly after ViviRen induction and tumor volume eventually decreased in the mLumiOpto (i.e., mAb-Exo-AAV+ViviRen) treatment group. In contrast, tumors continued to grow rapidly in all control groups (Fig. 8e). H&E staining of the parafilm sectioned TNBC tumors, harvested on Days 8-9 (i.e., 4-5 days after the last ViviRen induction), revealed severe cell death and reduced tumor cell density in mLumiOpto-treated mice compared to controls (Fig. 8f). Importantly, there was no apparent damage or injury in normal organs of mLumiOpto-treated (Fig. 8g) or control (data not shown) mice. These results demonstrate that the mAb-Exo-AAV-delivered mLumiOpto has great potential for cancer treatment with high anti-tumor efficacy and minimal toxicity.

The anti-cancer effectiveness of mLumiOpto was further assessed in immunocompetent mouse models. Briefly, the EGFR^+^ mouse TNBC 4T1 xenografted BALB/cJ female mice were administrated with either saline or anti-EGFR mAb-Exo-AAV on Day 0, followed by daily ViviRen injection on Days 4-6. IVIS imaging conducted after the first ViviRen induction showed a robust luminescent response in the 4T1 xenograft (Fig. 9a), confirming the tumor-specific and functional NLuc expression. Similar to the results observed in MDA-MB-231 xenografts, the growth of 4T1 tumors was significantly inhibited in the mAb-Exo-AAV group compared to the control group (Fig. 9b). Notably, the wet weight of terminal tumors in the mAb-Exo-AAV-treated mice accounted for only 6-8% of the wet weight observed in the control group (Fig. 9c). Intriguingly, we observed a significant infiltration of CD11c^+^ dendritic cells (DC) and CD8^+^ T cells in the TME of mLumiOpto-treated mice, as compared to the saline control (Fig. 9d), suggesting an enhanced tumoral immunity. To further investigate the immune regulation effect of mAb-Exo-AAV carrying mLumiOpto, Luminex multiplex assay of tumor tissues was performed to titrate the chemokines and cytokines in TEM. Significant upregulation of TFN-γ, IL-1β, IL-2, IL-4, IL-13 and IL-12p70 was detected in TEM while the modulation of immune suppression, as indicated by markers of IL-17A and IL-23, was minimal (Fig. 9e). Additional analysis of immunosuppressive markers CD33 and CD39 revealed slight downregulation of Tregs, 14.2-16.9% in saline group vs. 7.9-15.1% in mAb-Exo-AAV group, and minimal change of myeloid-derived suppressor cells (MDSCs), 7.3-11.6% in saline group vs. 8.39-11.3% in mAb-Exo-AAV treatment group (Fig. S9).

**Figure 9.**
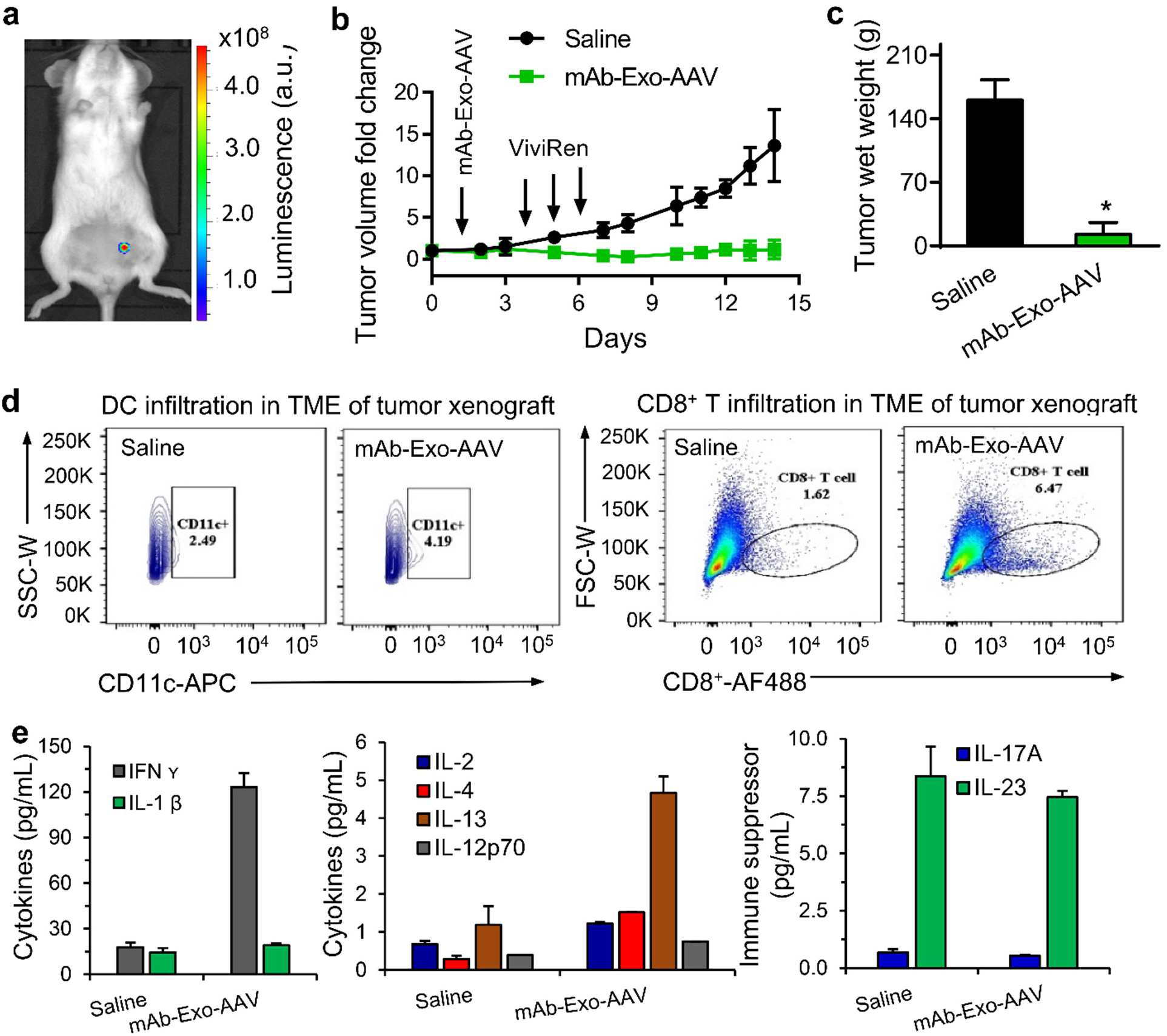
Evaluation of anti-cancer efficacy of mAb-Exo-AAV-delivered mLumiOpto in mouse 4T1 immunocompetent xenograft mouse model. (**a**) ViviRen induction triggered strong NLuc bioluminescent responses in mAb-Exo-AAV mice. (**b**) Prolonged ViviRen administration effectively inhibited tumor growth in mAb-Exo-AAV mice compared to control groups (saline). (**c**) The wet weight of terminal tumors in mAb-Exo-AAV mice was significantly lower than those of control (saline) mice. (**d**) Flow cytometry analysis showed enriched CD11C^+^ DC (left) and CD8^+^ T (right) cells in the tumor microenvironment (TME) of mLumiOpto (mAb-Exo-AAV+ViviRen)-treated mice compared with control (saline) mice. \**P*<0.05 vs. saline. n=6/group. (**e**) Luminex assay of tumor tissues showed upregulation of several cytokines, such as TFN-γ, IL-1β, IL-2, IL-4, IL-13 and IL-12p70, and downregulation of indicated downregulation of IL-17A and IL-23 by mAb-Exo-AAV.

## Discussion

Mitochondria has been considered as a potential therapeutic target for cancer treatment, but translating this concept into clinical practice has proven challenging. Although certain drugs targeting mitochondria have demonstrated promise in preclinical studies, their effectiveness in human clinical trials has been limited due to the low treatment efficacy, development of drug resistance in cancer cells and the lack of specificity leading to risk of side-effects. To overcome these limitations, we introduce a novel therapeutic strategy, mLumiOpto, that can specifically and directly destroy cancer mitochondria to induce cancer cell death. We utilize the AAV and mAb-Exo-AAV platform for the targeted delivery of mLumiOpto genes to cancer cells *in vivo* to maximize potency and minimize off-target effects. Our preclinical mouse xenograft models demonstrate that AAV (i.c.v. administration) and mAb-Exo-AAV (i.v. administration)-delivered mLumiOpto is effective at killing cancer cells and inhibiting tumor growth without causing noticeable side effects, highlighting its potential as a valuable tool for cancer research and targeted cancer therapy.

In recent years, optogenetics has revolutionized the precise and remote manipulation of cell membrane excitability, but its application to control intracellular organelles, particularly mitochondria, has remained limited. In this study, we expanded the application of optogenetics to damage cancer mitochondria and eliminate cancer cells. In particular, we introduced a novel approach called mitochondrial-targeted luminoptogenetics (Fig. 10), which enables specific and dynamic manipulation of mitochondria both *in vitro* and *in vivo*. Our strategy involved integrating the endogenous bioluminescence emitted by the NLuc-ViviRen pair into our previously developed mOpto system, resulting in an advanced external light-independent mLumiOpto technology. The rationale behind this design stems from the fact that luciferases emit light at specific wavelengths when coupled with suitable chemical substrates like luciferin (39–41). For instance, RLuc can emit blue light with λ_Peak_ of ∼470 nm in the presence of coelenterazine (42). Exploiting the spectral overlap between luciferase emission and rhodopsin absorption excitation, bioluminescence has proven successful in activating rhodopsin channels and controlling neuronal activity in live animals (43–45). In this study, we sought to identify the ideal luciferase that meets certain criteria, including monomeric structure, structural stability, high and sustained luminescence generation at low substrate doses, and non-toxicity to cells, to achieve dynamic and efficient mitochondrial control *in vivo*. Among several commonly used blue light-emitting luciferases and their variants, we determined that NLuc is the most suitable candidate for mLumiOpto technology. NLuc not only emits bright and sustained bioluminescence, even at low ViviRen concentrations, but it is also small in size (encoded by a 513-bp gene), ATP-independent, non-toxic to cells, and exhibits uniform intracellular distribution (28). Our *in vitro* studies demonstrate that NLuc-ViviRen pair-generated intracellular bioluminescence is sufficient and effective to activate the mitochondrial CoChR channel, resulting in ViviRen dose-dependent ΔΨ_m_ depolarization in the absence of external light stimulation. By harnessing the endogenous bioluminescence, mLumiOpto overcomes the technical challenges associated with delivering external light to deep tissues within the body and minimizes potential side effects on surrounding healthy tissues. The ability to manipulate mitochondrial function in freely-moving animals renders mLumiOpto a potent and versatile tool for *in vivo* investigations, ranging from elucidating the mechanistic underpinnings of mitochondrial dysfunction in various diseases to facilitating the development of mitochondrial-targeted therapeutic interventions.

**Figure 10.**
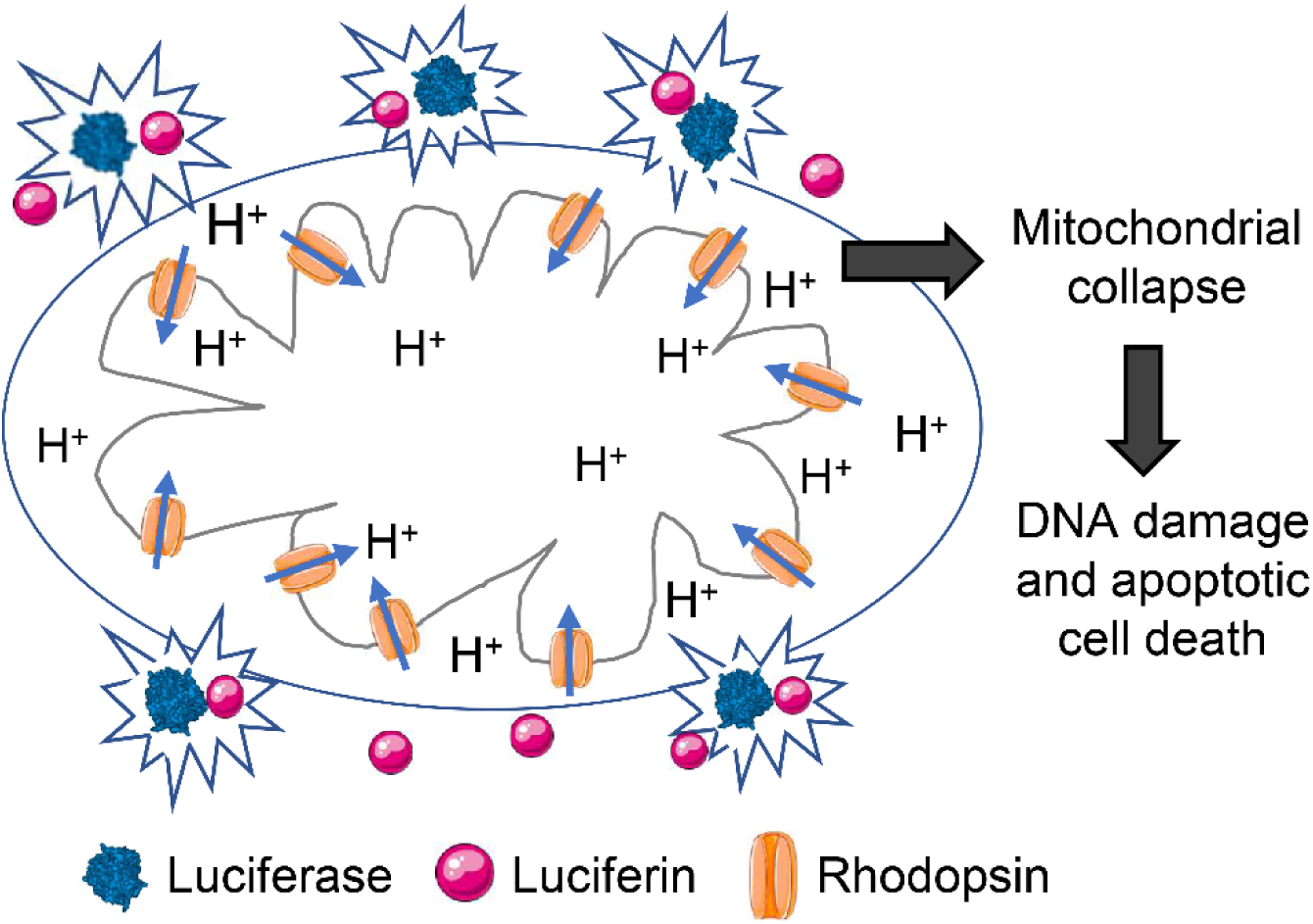
Diagram of mitochondria-targeted mLumiOpto technology. The blue light-gated rhodopsin channel protein (i.e., CoChR) and the emission spectra-matched luciferase (i.e., NLuc) are co-expressed within the same cancer cell, with CoChR in the inner mitochondrial membrane (IMM) and NLuc in the cytosol. The luciferin (ViviRen, an engineered NLuc substrate) induction prompts the emission of endogenous blue bioluminescence, activating the nearby cationic rhodopsin channels within the IMM. This activation leads to mitochondrial depolarization, subsequently causing damage to cancer mitochondria, induction of apoptosis, and instigation of DNA damage.

The capability of mLumiOpto in inducing cancer cell death was validated using *in vitro* cell lines, *ex vivo* patient tumor surrogates, and *in vivo* tumor xenograft mouse models. Consistent with our previous observations that mOpto mediated light intensity-dependent cytotoxicity, we found that mLumiOpto induced cancer cell death in a ViviRen dose-dependent manner. Importantly, mLumiOpto was cytotoxic to all treated cancer cell types. Dose dependence is vital for optimizing treatment efficacy while minimizing side effects, as different cancer cells may exhibit varying sensitivities to mLumiOpto-mediated cytotoxicity. Determining the optimal mLumiOpto dosing strategy may enhance treatment effectiveness further. Our animal studies further highlighted the ability of mLumiOpto to eliminate tumor cells *in vivo*. Treatment with mLumiOpto effectively inhibited tumor growth in various xenograft mouse models, including multiple TNBC subtypes and intracranial GBM. Our findings are particularly significant, as there is still a lack of effective treatment options for those highly aggressive and recurrent cancers. It is worth noting that neither ViviRen nor mLumiOpto expression alone exhibited any deleterious effects on cancer mitochondria and cell viability. Altogether, these findings indicate the robust and versatile nature of mLumiOpto as a promising approach for targeted cancer cell death in both preclinical and translational settings. Its ability to effectively combat tumor growth across diverse cancer types further positions mLumiOpto as a valuable therapeutic strategy for various malignancies.

Our studies provide valuable insights into the underlying mechanisms of mLumiOpto-induced cancer cell death. We found that this cell death primarily occurs through the intrinsic apoptotic pathway, accompanied by DNA damage, and is independent of mtROS and mPTP opening. These findings are of great significance as they not only confirm the functional expression of heterologous CoChR channels in IMM but also underscore mLumiOpto’s ability to eliminate cancer cells without relying on endogenous cancer types-associated proteins or signaling pathways. One of the major challenges in current cancer treatment is the development of drug resistance, which greatly comprises the efficacy of conventional chemotherapy, biological therapies, and other treatment modalities. Multiple mechanisms underlying drug resistance have been proposed, including cancer heterogeneity, the TME, cancer stem cells, inhibition of cell death pathways, and mutation or alterations in therapeutic targets (46,47). mLumiOpto employs heterologous genes that don’t target or rely on any cancer-associated proteins or pathways, such as EGFR, VEGF, ALK, PI3K/AKT/mTOR or p53, which are often impaired during long-term treatment. Consequently, mLumiOpto-based approaches may be less prone to develop drug resistance and more efficient in eliminating heterogeneous cancer cells compared with conventional therapies. These characteristics render mLumiOpto a promising strategy for overcoming the limitations associated with drug resistance and offer the potential for improved therapeutic outcomes in the treatment of diverse cancer types although a full evaluation is needed in future studies.

AAV vectors have gained considerable attention as a safe and promising approach for delivering therapeutic genes (48). The FDA has approved AAV-delivered Luxturna and Zolgensma to treat rare inherited blindness and spinal muscular atrophy, respectively, and numerous clinical studies exploring the application of AAV-based gene therapy for a wide range of diseases are currently underway (49–52). Despite the great potential, traditional free AAV-based gene therapies have several major challenges in cancer treatment, such as lack of cancer-specific targeting, low infection efficiency, pre-existing AAV neutralizing antibodies in a significant portion (∼30-66%) of the human population (53–56), and *de novo* production of anti-AAV antibodies by memory B lymphocytes (57,58). In this context, exosomes, the extracellular nanovesicles secreted by cells, have emerged as a promising vehicle for delivering therapeutic agents, including AAV vectors, due to their low antigenicity and toxicity. Compared with free AAV, Exo-AAV can help to protect the genetic material from degradation and increase its circulation stability. Additionally, mAbs can bind to specific receptors on the surface of cancer or tumor cells, directing the exosomes carrying therapeutic genes to the desired cells. Thus, the combination of mAbs and Exo-AAV (i.e., mAb-Exo-AAV) allows for targeted gene delivery, increasing the efficiency and specificity of mLumiOpto therapy. Several preclinical studies have demonstrated the potential of Exo-AAV for gene delivery to treat cancer such as Hemophilia A and B (59), liver disease (53), or lysosomal storage disorder by targeting the central nervous system (60). In this study, we established a novel platform of mAb-Exo-AAV by surface tagging a cancer surface receptor-targeting mAb (i.e., anti-EGFR mAb targeting 52-89% of TNBC) to Exo-AAV *via* DMPE-PEG-NHS linker as we previously described (61). mPEG-DSPE was also integrated into exosome membrane to further improve circulation stability. Our animal results demonstrated that mAb-Exo-AAV effectively targets tumors, achieving high-level and functional mLumiOpto expression specifically in tumor tissues with undetectable non-specific distribution in normal organs. It is worth noting that our strategy allows for facile surface conjugation of various mAbs, including dual mAbs, to Exo-AAV, thereby expanding the applicability of this approach to a broader range of patients and various cancer types or subtypes.

Remarkably, our study not only demonstrated the highly efficient and cancer-specific gene delivery capability of anti-EGFR mAb-Exo-AAV but also revealed its intriguing ability to induce an anti-tumor immune response in immunocompetent TNBC xenograft mouse models. The mechanisms underlying this enhanced tumoral immunity are multifaceted. First, it has been reported previously that EGFR activation plays a significant role in stimulating cancer proliferation, DNA damage repair, and metastasis (62). Clinical data further indicated that anti-EGFR mAbs (such as cetuximab or panitumumab) mediate antibody-dependent cell cytotoxicity within the intratumoral space and inhibit DNA repair *via* BRCA (63). Our mAb-Exo-AAV has a high mAb surface tagging rate with an optimal mAb:Exo-AAV ratio, which could facilitate the induction of immune responses within the TME. Second, the immune response triggered by AAV capsid immunity can reactivate memory CD8^+^ T lymphocytes through histocompatibility (MHC) class I presentation, leading to tumor destruction (57,64). After specifically targeting tumor cells, mAb-Exo-AAV is internalized to release free AAV intracellularly. The free AAV enters the nucleus, while its capsid undergoes proteasomal degradation, resulting in enhanced MHC surface expression, CD8^+^ T cell activation, and the induction of adaptive immunity within the TME. Third, in addition to T-cell infiltration, our flow cytometry analysis of freshly harvested tumor tissues revealed an increase in DCs. This observation aligns with the previously reported “cross-priming” mechanism (65), whereby apoptotic cancer cells release tumor antigens into the TME. These antigens are subsequently captured by antigen-presenting DC cells, facilitating the activation of CD8^+^ T cells. Activated CD8^+^ T cells within the TME can selectively target cancer cells through T-cell receptors, effectively eliminating them. Consequently, the elimination of cancer cells creates a more favorable TME that fosters the infiltration of immune cells, further boosting tumoral immunity. With that said, multiple immunocompetent models and humanized mouse models will need to be used to fully investigate the immune effects in the TME, decipher the underlying mechanisms, and examine the potential synergistic anti-cancer efficacy of mLumiOpto technology and the gene delivery vehicle mAb-Exo-AAV in future studies.

In summary, our study introduces mLumiOpto, an innovative mitochondrial-targeted luminoptogenetic approach that enables dynamic manipulation of cancer mitochondria and triggers cytotoxicity. Our findings highlight the potential of mAb-Exo-AAV-delivered mLumiOpto as a promising strategy for targeted induction of cancer cell death and activation of immune response in the TME with specificity. These advancements provide valuable insights for the development of novel therapeutic strategies that can address major challenges in cancer treatment, including reduced drug resistance and enhanced efficacy. Additional evaluations using more clinically relevant animal models, such as metastatic models and PDX xenograft humanized mouse models, are essential to further explore the translational potential of this promising therapeutic strategy. Furthermore, the comprehensive assessments of Investigational New Drug (IND)-directed toxicology, biodistribution, pharmacokinetics and pharmacodynamics are essential for laying the foundation for future clinical trials. Importantly, besides GBM and TNBC, mLumiOpto technology has great potential for treating other challenging cancers, such as recurrent GBM and non-small cell lung cancers, by substituting the cancer-targeting mAb on the surface of Exo-AAV. Finally, mitochondria play an essential role in tumorigenesis, metastasis, and stemness, so its capability of dynamically depolarizing mitochondria renders mLumiOpto a powerful tool in mechanistic studies.

## METHODS

Five-week-old nude (J:NU HOM Homozygous for Foxn1 <NU>) mice, NSG (NOD.Cg-Prkdc^scid^ Il2rg^tm1Wjl^/SzJ) female mice, and BALB/cJ female mice were purchased from Jackson Lab. After one week of acclimatization, mice were used to generate tumor xenograft models for *in vivo* evaluation of mLumOpto technology. All animal studies conformed to the Laboratory Animals Guideline of the US National Institutes of Health (Publication No. 85-23). The animal protocols were approved by the Institutional Animal Care and Use Committee at Ohio State University (IACUC-2022A00000035 and IACUC-2022A00000029).

### Cell lines, seed cultures and media

Viral Production Cells 2.0 (VPC, Gibco, Buffalo, NY) were used to produce AAV and Exo-AAV. The cervical cancer cell line HeLa (ATCC) was used for the general characterization of mLumiOpto technology. The human TNBC cell lines MDA-MB-231 (GenTarget, San Diego, CA), MDA-MB-468 (GenTarget) and BT-20 (ATCC), and GBM cell lines U251 (MilliporeSigma, Burlington, MA), drug-resistant U251-TMZ (in-house developed), U87 and GL261 (Creative Bioarray, Shirley, NY) were used for *in vitro* cytotoxicity, surface binding, AAV gene delivery, gene expression, or mechanism studies. The MDA-MB-231-FLuc (GenTarget), 4T1-FLuc (ATCC), U87, U251, and U87-FLuc (ATCC) were used to develop xenograft mouse models (immunocompromised and immunocompetent) for evaluating the *in vivo* anti-tumor efficacy of mLumiOpto.

All cell culture media, supplements and other reagents were purchased from Gibco unless otherwise specified. HEK 293A cells were cultured in Dulbecco’s Modified Eagle Medium (DMEM) supplemented with 10% FBS, 2 mM L-glutamine, 100 U/mL Penicillin/100 µg/mL Streptomycin, and MEM Non-Essential Amino Acids in T-flasks. The seed culture of VPC was maintained in a chemically defined viral production medium supplemented with 4 mM GlutaMAX in shaker flasks on an orbital shaker at 135 rpm. HeLa cells were cultured in DMEM supplemented with 10% (v/v) Fetal Bovine Serum (FBS) and 2 mM L-glutamine in T25 or T75 flasks. The MDA-MB-231, MDA-MB-468 and BT-20 cells were cultured in DMEM/F12 medium supplemented with 10% FBS, 4 g/L glucose, 4 mM L-glutamine, 100 U/mL Penicillin and 100 µg/mL Streptomycin in T-flasks. U251 and U251-TMZ cells were maintained in EMEM with 2 mM L-glutamine, 1% non-essential amino acids (NEAA), 1 mM sodium pyruvate and 10% FBS in T25 or T75 flasks. U87 cells were maintained in EMEM with 10% FBS and 8 µg/mL Blasticidin. All cell lines were maintained between 10% and 80% confluence and kept at 37 °C with 5% CO_2_ (8% CO_2_ for VPC cells) in a humidified CO_2_ incubator (Eppendorf, Enfield, CT). The viable cell density (VCD) and viability were measured using TC20 automated cell counter (Bio-Rad, Hercules, CA) or hemocytometer and trypan blue (Fisher Scientific, Hanover Park, IL).

### Plasmid construction

#### CMV-ABCB-CoChR-eYFP

The ABCB, CoChR and eYFP gene fragments were amplified from CAG-ABCB-ChR2-eYFP(25), AAV-Syn-CoChR-GFP (Addgene #59090)(66), and pcDNA3.1-PsChR2-eYFP (Addgene #69057) (67), respectively. The PCR primers are ABCB-forward, ABCB-reverse, CoChR-forward, CoChR-reverse, eYFP_1-forward, and eYFP_1-reverse (Supplemental Table 1). These gene fragments were cloned into pcDNA3.1-PsChR2-eYFP backbone vector using the HiFi Assembly Kit (New England Biolabs, lpswich, MA).

#### CMV-ABCB-CoChR-mCherry

The ABCB-CoChR gene fragments were amplified from CMV-ABCB-CoChR-eYFP and cloned into pcDNA3.0-Magneto2.0-p2A-mCherry (Addgene #74308) backbone vector using the HiFi Assembly Kit. The primers ABCB-CoChR_1-forward and ABCB-CoChR_1-reverse are listed in Supplemental Table 1.

#### CMV-NLuc-2A-ABCB-CoChR-mCherry

The NLuc, 2A, and ABCB-CoChR-mCherry gene fragments were PCR amplified from pNL-CMV-NLuc (Promega #N1091), pcDNA3.0-Magneto2.0-p2A-mCherry, and CMV-ABCB-CoChR-mCherry, respectively. The amplified genes were cloned into the CMV-ABCB-CoChR-mCherry vector using the HiFi Assembly Kit. The PCR primers are NLuc_1-forward, NLuc_1-reverse, 2A-forward, 2A-reserve, ABCB-CoChR_2-forward, and ABCB-CoChR_2-reserve (Supplemental Table 1).

#### CMV-NLuc-GFP-2A-ABCB-CoChR-mCherry

The NLuc, GFP, and 2A-ABCB-CoChR-mCherry gene fragments were PCR amplified from pNL-CMV-NLuc, CMV-myc-mito-GFP, and CMV-NLuc-2A-ABCB-CoChR-mCherry, respectively. The amplified genes were cloned into CMV-NLuc-2A-ABCB-CoChR-mCherry vector using the HiFi Assembly Kit. The primers used are NLuc_2-forward, NLuc_2-reserve, GFP-forward, GFP-reserve, 2A-ABCB-CoChR-forward, and 2A-ABCB-CoChR-reserve (Supplemental Table 1).

#### pAAV-D/J8-cfos-NLuc-2A-ABCB-CoChR

The NLuc-2A-ABCB-CoChR gene fragment was PCR amplified from CMV-NLuc-2A-ABCB-CoChR-mCherry, and cloned into the pAAV-D/J8 AAV expression vector (Cell Biolabs, San Diego, CA) following the manufacture instruction. The primers are NLuc-CoChR_2-forward and NLuc-CoChR_2-reverse (Supplemental Table 1). The sequence of the construction was confirmed with sequencing primers forward: 5’-GGATTGACGGGAACAG-3’, and reverse: 5’-GGTGTATATCGAGAGC-3’.

### AAV and Exo-AAV production and purification

#### Production

The small-scale (30 or 60 mL) productions of AAV and Exo-AAV were performed in 125 or 250-mL shaker flasks using VPC 2.0 in viral production medium at 37 °C, 135 rpm and 8% CO_2_. The large-scale production was performed in 2-L stirred-tank bioreactor (Disteck, Cedar Falls, IA) at 37 °C, pH 7.0, 210 rpm and DO 40%. VPC cells were co-transfected at VCD of 3×10^6^ cells/mL with three plasmids, i.e., AAV-D/J8-cfos-NLuc-2A-ABCB-CoChR, AAV-DJ/8 Rep-Cap and AAV-D/J8 Helper (1:3:1), at plasmid DNA:cell ratio of 0.5 µg:10^6^ cells. The formulation of transfection mixture per liter of culture volume was 1.5 mg total plasmid DNA, 10% (v/v) viral-plex complexation buffer, 0.6% AAV-MAX transfection reagent, and 0.3% AAV-MAX transfection booster. The enhancer (1%) was added to the VPC culture immediately before plasmid transfection. Production culture was sampled daily to monitor cell growth using TC20 automated cell counter, glucose level using Glucose 201 DM System (HemoCure, Brea, CA), AAV titer using qRT-PCR, and Exo-AAV titer using nanoparticle tracking analysis (NanoSight, Malvern Panalytical, Malvern, UK) (68), respectively. VPC cells containing AAV were harvested at 72 hrs post-transfection for AAV isolation, and the spent medium containing Exo-AAV was collected when viability dropped to 50% for Exo-AAV purification.

#### Clarification

In the end of AAV/Exo-AAV production, culture broth was centrifuged at 4 °C and 1,000 x g for 20 minutes. The supernatant containing raw Exo-AAV was collected and clarified using Cytiva Supracap 50 depth filter capsules with dual-layer media grade PDK5 (1.5-20 µm) and PDE2 (0.2-3.5 µm) with regenerated cellulose membrane, followed with 100 kDa MWCO regenerated cellulose membrane for size-exclusion primary purification with buffer exchange and concentration for further Exo-AAV purification. VPC cell pellet was re-suspended in 10% AAV-MAX lysis buffer (Gibco), followed by three freeze-thaw cycles (incubation in ethanol/dry ice bath for 30 minutes and in 37 °C water bath for 15 minutes), and further incubated with 2 mM MgCl_2_ and 90 U/mL benzonase (MilliporeSigma, Burlington, MA) at 37 °C for 60 minutes. After cell lysis was confirmed with microscopy, the lysate was clarified by centrifugation at 4 °C and 4,500 × g for 30 minutes, and the supernatant was filtered using 0.2-µm PES membrane to remove cell debris. The filtrate was partially purified, preconditioned, and concentrated using Vivaspin Turbo column MWCO 100 kDa (Sartorius, Bohemia, NY) for further AAV purification.

#### Purification

NGC liquid chromatography (Bio-Rad) equipped with a 4.7-mL Capto Core 400 size exclusion column (Cytiva, Marlborough, MA) was used to purify Exo-AAV. Briefly, the column was washed with equilibration buffer (formulation of 50 mM Tris, 50 mM NaCl, pH 7.3), loaded with clarified Exo-AAV sample, and eluted with buffer of 50 mM Tris, 1.2 M NaCl, pH 7.3 at a flow rate of 1.3 mL/minute. The purified Exo-AAV was processed with dialysis for buffer exchange and a refrigerated vacuum concentrator (Fisher Scientific) for concentration. NGC liquid chromatography, equipped with a 5-mL Foresight Nuvia HPQ anion-exchange column prepacked with CHT ceramic hydroxyapatite XT mixed-mode chromatography media (Bio-Rad) or 25-mL column in-house packed with the same CHT media, was used to purify free AAV. HPQ column was equilibrated with buffer A (25 mM Tris-HCl, 20 mM NaCl, pH 8.0), loaded with the buffer A-preconditioned AAV sample, and eluted step wisely using buffers A and B (25 mM Tris-HCl, 1 M NaOH, pH 8.0) at a flow rate of 0.6 mL/minute. The primarily purified AAV was further proceeded with ultrafiltration using 100-kDa MWCO PES membrane column to remove small protein impurities, desalt, exchange with formulation buffer (1x PBS, 5% Sorbitol, and 350 mmol/L NaCl), and concentrate as previously described (38).

### mAb-Exo-AAV construction and titration

#### mAb-Exo-AAV construction

The biosimilar of Cetuximab, anti-EGFR mAb (Bio X Cell, Lebanon, NH), was tagged to the surface of Exo-AAV *via* mPEG-DSPE linker to generate the TNBC-targeting mAb-Exo-AAV. Following the procedure developed in our previous study (61,69,70), Exo-AAV was labeled with fluorescent dye Cy5.5 PE (only for IVIS imaging) and modified by mPEG-DSPE with a molar ratio of 1:10,000:6,000,000 (Exo-AAV:Cy5.5:mPEG-DSPE). The Exo-AAV-PEG-Cy5.5 was conjugated with anti-EGFR mAb *via* DSPE-PEG-NHS linker with a molar ratio of 1:2,680:13,000.

#### Titration and characterizations

The purified AAV was digested with DNAse I to extract ssDNA and titrated using RT-PCR with primers (forward: 5’-ATTGTCCTGAGCGGTGAAA-3’, reverse: 5’-CACAGGGTACACCACCTTAAA-3’). The size distribution, morphology, biomarkers and purity of mAb-Exo-AAV were characterized using NanoSight, transmission electron microscope (TEM), and Western blotting. The AAV packed in each exosome was titrated using the same primers and Exo-AAV was titrated using nanoparticle tracking analysis (NanoSight) to calculate AAV packing rate in Exo (i.e., AAV copy per Exo particle).

### Flow cytometry analysis

The mAb-EV-AAV samples were labelled with fluorescent dye Cy5.5 to test the TNBC cell surface binding rate. Briefly, 1×10^12^ ptc of mAb-Exo-AAV was mixed with 16.7 nmole of Cy5.5, shaking incubated overnight in dark at room temperature, and purified using Vivaspin 100 kDa column (Fisher) to remove free dye. Then 1×10^6^ of MDA-MB-231 or MDA-MB-468 cells were stained with 4×10^10^ ptc of mAb-Exo-AAV at room temperature for 30 mins and washed with PBS twice. The stained cells were applied to BD LSRII flow cytometer (BD Biosciences, San Jose, CA, USA) and data were analyzed with FlowJo V5.0 (TreeStar, Inc., Ashland, OR, USA). Gating was set where negative samples (stained with mAb-Exo-AAV) have <0.5% fluorescent population.

### Confocal imaging

#### Colocalization analysis

Cells cultivated on a 15-mm glass-bottom dish were transfected with *CMV-ABCB-CoChR-eYFP* plasmid. Forty-eight hours after transfection, cells were loaded with MitoTracker Deep Red (250 nM) for 30 minutes. The localization of CoChR-YFP and MitoTracker was simultaneously imaged with 543 nm argon laser and 635 nm laser diode lines using Olympus FV1000 confocal microscope (Olympus America, Center Valley, PA). For NLuc and CoChR co-expression analysis, cells were transfected with CMV-NLuc-GFP-2A-ABCB-CoChR-mCherry plasmid. Forty-eight hours later, the expression of NLuc-GFP and CoChR-mCherry was imaged with the 488 and 545 laser lines, respectively. For colocalization analysis, the confocal images were processed offline using ImageJ software (National Institutes of Health, Bethesda, MD). The overlapping was quantified by calculating the Manders coefficient.

#### Mitochondrial depolarization and ΔΨ_m_ measurement

The mLumiOpto-treated or control cells were stained with fluorescent mitochondrial membrane potential dye TMRM (100 nM) or Mitoview 633 (25 nM). The fluorescence of TMRM and Mitoview 633 was imaged with the 543 nm and 635 nm laser, respectively, and analyzed using ImageJ software.

#### Analysis of AAV or mAb-Exo-AAV transduction in vitro

As described in our previous studies (71,72), AAV or mAb-Exo-AAV carrying *cfos-NLuc-2A-ABCB-CoChR* genes was labelled with fluorescent dye Cy5.5 or Cy7 and incubated with TNBC MDA-MB-468 cells that were stained with DAPI for 20 minutes. The transduction of AAV-Cy5.5 was detected with an Olympus FV1000 confocal microscope. The nucleus stained with DAPI was imaged with the 405 nm laser line and Cy5.5 was imaged with the 640 nm laser line, respectively.

### *In vitro* cytotoxicity assay

Cancer (TNBC or GBM) cells were seeded onto 96-well plates at a density of 5×10^4^ cells/mL, and incubated with mLumiOpto plasmid (DNA:cells=1.2 µg:1×10^6^ cells) or AAV (MOI of 100,000) at 37 °C and 5% CO_2_ in the incubator for 48 hours. Then ViviRen (0-60 µm) was added to the culture in well plates. Two days later, cell viability was measured using CellTiter-Glo Luminescent Cell Viability Assay (Promega, Madison, MI), as previously described(72).

To delineate the mechanistic pathway underlying mLumiOpto-mediated cytotoxicity, cells were treated with mLumiOpto for 48 hours with or without the presence of 20 µM Z-VAD-FMK (a pan-caspase inhibitor), 100 µM 7-Cl-O-Nec-1 (a necroptosis inhibitor), 10 µM bafilomycin A (BfA, an autophagy inhibitor), 10 µM cyclosporin A (CsA, an mPTP opening inhibitor), 100 µM Z-DEVD-FMK (a caspase-9 specific inhibitor), 20 µM Z-LEHD-FMK (a caspase-3 specific inhibitor), 20 µM Z-IETD-FMK (a caspase-8 specific inhibitor), or 300 nM MitoQ (a mitochondrial-specific antioxidant). Cell viability was measured using a Trypan Blue exclusion assay or TC20 automated cell counter (BioRad, Hercules, CA).

### Bioluminescence imaging

#### In vitro imaging

Cells were seeded in clear-bottom black well-plates (Corning, Corning, NY) and cultured with mLumiOpto plasmid or mAb-Exo-AAV. Forty-eight hours later, the culture medium was replaced with a colorless culture medium containing a varying dose of ViviRen (0-30 µM). The bioluminescence was detected at 0, 2, 4, 6 and 20 hours using IVIS Lumina Series III (PerkinElmer, Waltham, MA) at 470 nm.

#### In vivo imaging

TNBC tumor-bearing mice were injected with a single dose of mLumiOpto (1×10^10^ ptc/g-BW of mAb-Exo-AAV *via* intravenous injection) and ViviRen (2 μg/g-BW *via* intravenous injection). Twenty-four hours later, mice were imaged with IVIS Lumina Series III to measure the NLuc luminescence. To evaluate the tumor-specific targeting and biodistribution of mLumiOpto, the TNBC MDA-MB-231-FLuc xenograft mice were injected with 1×10^10^ ptc/g of anti-EGFR mAb-Exo-AAV *via* tail vein. Twenty-four hours later, mice were imaged under IVIS with an exposure time of 10 seconds. The tumor-specific targeting was determined by: 1) analyzing the overlay of NLuc luminescence (mAb-Exo-AAV) and tumor in live-animal IVIS imaging, as previously described (38), 2) *ex vivo* IVIS imaging of the harvested major organs (brain, heart, lung, liver, kidney, spleen) after mice were sacrificed, and 3) transcript analysis of tumor and organs (heart, brain, lung, kidney, intestine, spleen, pancreas, stomach, colon, liver) using qRT-PCR.

### Magnetic Resonance Imaging (MRI) imaging

The Magnetic Resonance Imaging (MRI) imaging was performed with the BioSpect 94/30USR system (Bruker BioSpin; Billerica, MA) and ParaVision 6.0 software provided by the Ohio State University Small Animal Imaging Core Facility. T2-weighted scans were acquired with flowing parameters of TR/TE: 2500/33 (ms), FA: 180 (degree), NEX: 2, FOV: 20 mm*15.313, matrix: 256*196, 1 mm slice thickness, 1 mm slice distance and 18 slices. Mice were anesthetized and 0.2 mmole/kg Gadolinium-based contrast agent was administrated intraperitoneally before imaging. Then the mice were secured on an animal bed and placed in the MRI scanner for imaging. A rectal thermometer with body contact was used to measure the body temperature. The respiration and heart rate of mice were monitored using the Small Animal Monitoring System (Model 1025, Small Animals Instruments, Inc. Stony Brook, NY) during the imaging session.

### Echocardiography

Transthoracic echocardiography was performed on mice to evaluate the cardiac function of mice using a 30 MHz RMV707B transducer and VisualSonics Vevo3100 High-Resolution System (VisualSonics, Toronto, ON, Canada) as previously described (73,74). In brief, mice were anesthetized with 1.5% inhaled isoflurane anesthesia (with 100% supplemental oxygen) and placed on a heated, bench-mounted adjustable rail system. The two-dimensionally directed M-mode images were captured from the long-axis views. Left ventricular (LV) end-diastolic and end-systolic dimensions, and LV end-diastolic posterior wall thickness, were measured from the M-mode images. LV fractional shortening was calculated with VisualSonics V1.3.8 software.

### Xenograft mouse models

#### Human GBM xenograft model

Six-week-old nude (J:NU HOM Homozygous for Foxn1 <NU>) mice, with an equal number of males and females, were stereotactically injected with human GBM cells(69,71). Briefly, 0.5×10^5^ U87 cells were suspended in 3-µL growth medium and implanted into the frontal region of the cerebral cortex at a rate of 0.4 µL/min using Stoelting Just for Mouse Stereotaxic Instrument (Stoelting, Wood Dale, IL). The burr hole in the skull was closed with sterile bone wax and 5 mg/kg of carprofen was provided immediately before surgery and every 12-24 hours for 48 hours post-surgery. The intracranially xenografted mice were monitored daily for one week and randomized into four groups (n=10/group). Then mice were treated with saline (control), AAV only (0.5×10^11^ vg, control), mLumiOpto at AAV doses of 0.5×10^11^ vg and 1.0×10^11^ vg on Days 7 and 14 *via* i.c.v. injection, followed by three s.c. injections of ViviRen (37 µg) at 3 days post AAV administration in mLumiOpto treatment groups.

GBM PDX line was provided by Dr. Jann Sarkaria at Mayo Clinic and maintained at low passages (2–4) in NSG mice following our established protocol(69,71). Fresh frozen PDX tissues were thawed, minced into small fragments (∼1 mm^3^), and subcutaneously (s.c.) implanted into NSG mice. The PDX xenograft mice were randomized into two groups (n=5), treated with AAV (1×10^13^ vg/kg) carrying mLumiOpto genes *via* tail vein with 5 injections from Day 4 to day 8, and followed with four s.c. injections of ViviRen (37 µg) from Day 13 to Day 16. Tumor volume was measured by a caliper and body weight was monitored every two or three days for 35 days.

#### Human TNBC xenograft model

Five million human TNBC MDA-MB-231-FLuc or MDA-MB-231 cells were orthotopically injected into the mammary fat pad of 6-week-old NSG (NOD.Cg-Prkdc <SCID> Il2rg<TM1WJL>/SzJ) female mice to construct immunocompromised models. When the tumor volume reached ∼75-100 mm^3^, mice were i.v. administrated with anti-EGFR mAb-Exo-AAV (2×10^10^ ptc/g), free AAV (2×10^10^ vg/g), ViviRen (2 μg/g), or saline in 50 µL through tail vein injection (n=6/group). Four days later, the mice in mAb-Exo-AAV group and ViviRen group received daily ViviRen injections (2 μg/g) for three consecutive days. Tumor volumes were measured using an external caliper and recorded every two days. At the end of treatment, tumor tissues and major organs were harvested for parafilm section, H&E staining, and biochemical analysis.

#### Mouse TNBC xenograft model

Three million mouse 4T1 cells were subcutaneously injected into the mammary fat pad of 6-week-old BALB/cJ female mice to construct immunocompetent models. Mice were randomly divided into two groups (n=6/group) when tumor volume reached ∼75-100 mm^3^, and received Saline or anti-EGFR mAb-Exo-AAV (2×10^10^ ptc/g) on Day 0 (i.v. injection), followed by ViviRen administration (2 μg/g) *via* tail vein injection on Days 4-6. Tumor volumes were measured every two days using an external caliper, and the wet weight of terminal tumors was measured at the end of the study. Tumor tissues were used for flow cytometry analysis to detect immune cells infiltration in TME.

### Western blotting

Cells were added to ice-cold Pierce^TM^ RIPA lysis buffer (Thermo Fisher). Cell lysates were agitated gently at 4 °C for 1 hour, followed by centrifuge and supernatant collection. Protein concentration in the supernatants was determined using the BCA assay. Cell lysate or AAV samples were subjected to SDS-PAGE in NuPAGE 4-12% Bis-Tris gels. After electrophoresis, proteins were electro-transferred to a PVDF membrane. The blotted membrane was then blocked with TBS containing 5% fat-free milk powder and 0.1% Tween-20 for 1 hour at room temperature and incubated with specific primary antibodies overnight at 4 °C. The membrane was rinsed three times and incubated with HRP-conjugated secondary antibodies (anti-rabbit or mouse depending on the primary antibody, 1:3,000 for 1 hour at room temperature). The membrane was treated with Luminata Forte Western HRP substrate (MilliporeSigma, Burlington, MA) and then imaged with a MyECL imager (Thermo Fisher, Asheville, NC). Quantification analysis of blots was performed with ImageJ software. Targeted bands were normalized to GAPDH.

### RNA isolation and transcript expression analysis

Total RNA was extracted and purified from the cardiac tissues using the RNeasy mini kit (Qiagen, Germantown, MD). cDNA was synthesized from 500 ng of RNA using QuantiTect Reverse Transcription Kit (Qiagen). Quantitative real-time PCR was performed using Select Master Mix (Thermo Fisher, Asheville, NC) in a BioRad IQ5 detection system (Bio-Rad, Portland, ME). The transcript level of the CoChR gene was normalized to the average levels of Gapdh and Rpl32. PCR primers are listed in Supplemental Table 2, and their specificity was confirmed with 1% agarose gel electrophoresis and melt curves. The fold difference for the mRNA expression level was calculated using 2^−ΔΔCt^.

### Immunofluorescence

The TNBC (MDA-MB-231) cells cultivated on glass coverslips or tissue sections were fixed in 4% formaldehyde in PBS and then treated with PBS containing 10% goat serum and 0.3% Triton X-100 to block nonspecific staining. Cells were then incubated overnight at 4°C with anti-TOMM20 (1:200 dilution) and anti-cytochrome C (1:200 dilution) primary antibodies (Abcam, Waltham, MA). Thereafter, cells were stained with 1:200 diluted secondary antibodies labeled with AF488 and/or AF647 in 1% BSA, 1% goat serum, and 0.3% Triton X-100 in PBS. Finally, coverslips were mounted on slides and imaged using an Olympus FV1000 confocal microscope.

### Hematoxylin and eosin (H&E) staining

The tumor tissue and normal organs (heart, brain, lung, liver, spleen, kidney) were embedded in paraffin and sectioned at 5 μm. After deparaffinizing, the slides were stained with hematoxylin and eosin solution and imaged using the high-performance Nikon microscope (Irving, TX) as previously described (72).

### Tolerated dosage study

Varying doses of mAb-Exo-AAV (1, 2, 20, 40, and 80×10^13^ ptc/kg-BW) and ViviRen (2 mg/kg-BW) were administered intravenously into 6-week-old wild-type mice (BALB/cJ or C57BL/6J) (n=4/group). Mouse body weight was measured twice a week for 2 weeks post-injection. At the end of the study, mice were subjected to echocardiography to measure left ventricular contractile function. The blood samples were extracted by cardiac puncture for whole blood analysis using HemaVet 950FS (Drew Scientific, Miami Lakes, FL). The leukocytes (white blood cell, neutrophil, lymphocyte, monocyte, eosinophils, basophil), erythrocytes (red blood cell, hemoglobin, hematocrit, hepatitis, mean corpuscular hemoglobin), and thrombocytes (platelet) were counted. The major organs (brain, heart, lung, liver, spleen, kidney) were isolated, sectioned and used for H&E staining.

### Tumoral immunity analysis

#### Flow cytometry

The freshly isolated tumor tissues were dissociated with Tissue Dissociation/Single Cell Isolation Kit (101 Bio LLC, Sunnyvale, CA) following the manufacturing procedure. The dissociated tumor cells were stained with AF488 anti-CD8 antibody and APC anti-CD11c antibody (BioLegend, San Diego, CA) and subjected to flow cytometry analysis to access infiltrated immune cells in the tumor microenvironment. Gating was set where negative samples stained with unlabeled antibodies have <0.5% fluorescent population.

#### Luminex assay

The multiplexing kit EPX260-26088-901 (Thermo Fisher) was used to titrate tumor chemocytokines using the Luminex MAGPIX (Luminex Corporate, Austin, TX, USA) following the manufacturer’s instructions.

### Blood analysis

#### Complete blood count

The whole blood samples were drawn from the heart for blood cell count using HemaVet 950FS (Drew Scientific, Miami Lakes, FL, USA). The red blood cell (RBC), lymphocyte, hemoglobin, monocyte, white cell count (WBC), and neutrophil counts were compared and presented.

#### Serum chemical assay

About 10-50 µL of blood samples were taken from the tail and centrifuged at 2,000 g for 10 mins to collect serum. ELISA assays were conducted to titrate alanine transaminase (ALT, ab282882, Abcam), aspartate aminotransferase (AST, ab263882, Abcam), aspartate aminotransferase (BUN, EIABUN, Fisher), and creatinine (50-194-7694, Fisher) following biomanufacturer protocols.

### Antibodies and chemicals

Mouse primary antibodies and HRP-conjugated secondary anti-mouse antibodies were obtained from Abcam. TOMM20 and cytochrome c antibodies were purchased from Abcam and Cell Signaling Technology (Danvers, MA), respectively. MitoSox and MitoTracker were purchased from Life Technologies. PVDF membrane and T-PER tissue protein extraction reagents were obtained from Thermo Scientific (Waltham, MA). All other reagents were from MilliporeSigma.

### Statistical analysis

The experimental data were reported as mean ± standard error of the mean (SEM). Statistical comparisons among groups were performed using the two-way ANOVA Tukey’s multiple comparisons test or one-way ANOVA Holm-Sidak’s multiple comparisons test. *P*<0.05 was considered statistically significant. The distribution of data was tested using the Shapiro-Wilk normality test.

## Acknowledgments

This work was supported by DoD BCRP W81XWH2110066/67 (X.M.L. and L.Z.), NIH NCI 1R01CA262028-01A1 (X.M.L. and L.Z.), and NIH R01HL156581 (L.Z.). The authors thank the Comparative Pathology, Digital Imaging Shared Resource (CPDISR), Small Animal Imaging Shared Resource (SAISR), Flow Cytometry Shared Resource, Campus Microscopy & Imaging Facility (CMIF), and Center of Electron Microscopy and Analysis (CEMAS) at the Ohio State University for the assistance with tissue sectioning, IVIS imaging, flow cytometry, nanoparticle tracking assay, confocal imaging, and TEM imaging.

## Author contributions

L.Z. and X.M. L. conceived the idea, designed mLumiOpto technology, and supervised all experiments. P.E. developed mLumiOpto and performed *in vitro* evaluations. K.C., S.K., and T.V. produced, purified, titrated and characterized AAV, Exo-AAV and mAb-Exo-AAV. K.C. and Y.S. performed *in vivo* evaluations. S.K. performed Western blotting, qRT-PCR and immunofluorescence assay. K.C., P.E., X.M.L. and L.Z. analyzed data. M.R. edited the manuscript, and X.M.L. and L.Z. wrote and edited the manuscript.

