## Supplemental information for "An Innovative Mitochondrial-targeted Gene Therapy for Cancer Treatment"

**Short title**: Targeting Cancer Mitochondria *in vivo*

*Corresponding:
Lufang Zhou
460 W 12th Avenue, Columbus, Ohio, 43210, USA.


Xiaoguang "Margaret" Liu
151 West Woodruff Avenue, Columbus, OH 43210, USA


[+]These two authors have an equal contribution.

**List of Online Supplemental Table**

| Name | Sequence |
|---|---|
| ABCB-forward | TAAGCTTGGTACCGAGCTCGGATCCCACCATGCGCGCCCCTTC |
| ABCB-reverse | TTCCCAGCATAACTGCAGCTGACAGTCTCCC |
| CoChR-forward | AGCTGCAGTTATGCTGGGAAACGGCAGC |
| CoChR-reverse | TCACTAGCATTGCTACTACCGGTGCCGC |
| eYFP_1-forward | GGTAGTAGCAATGCTAGTGAGCAAGGGC |
| eYFP_1-reverse | ACCTTCGAACCGCGGGCCCTCTAGATTACTTGTACAGCTCGTCC |
| ABCB-CoChR_1-forward | ACCCAAGCTTGGTACCGGGTCTAGACACCATGCGCGCCCCTTC |
| ABCB-CoChR_1-reverse | CCTCCTCGCCCTTGCTCACGGATCCCACTGTCTCCTCGTCCTCCTG |
| NLuc_1-forward | ACTCACTATAGGGAGACCCACACCATGGTCTTCACACTCG |
| NLuc_1-reverse | TGCCTGATCCCGCCAGAATGCGTTCGCAC |
| 2A-forward | CATTCTGGCGGGATCAGGCAGCGGCGCC |
| 2A-reserve | GGGCGCGCATGGGACCGGGGTTTTCTTCCACG |
| ABCB-CoChR_2-forward | CCCCGGTCCCATGCGCGCCCCTTCTGCT |
| ABCB-CoChR_2-reserve | CCTCCTCGCCCTTGCTCACGTGCTACTACCGGTGCCGC |
| NLuc_2-forward | ACTCACTATAGGGAGACCCACACCATGGTCTTCACACTCG |
| NLuc_2-reserve | TGCTAGCCATCGCCAGAATGCGTTCGCAC |
| GFP-forward | CATTCTGGCGATGGCTAGCAAAGGAGAAG |
| GFP-reserve | CCCTGCCCTCGTAGAGCTCATCCATGCC |
| 2A-ABCB-CoChR-forward | TGAGCTCTACGAGGGCAGGGGAAGTCTTC |
| 2A-ABCB-CoChR-reserve | CCTCCTCGCCCTTGCTCACGCACTGTCTCCTCGTCCTC |
| NLuc-CoChR_1-forward | GTACCGGGTCTAGAGCCACCATGGCTTCCAAG |
| NLuc-CoChR_1-reverse | CCTTGCTCACCATTGCTACTACCGGTGC |
| eYFP_2-forward | GGTAGTAGCAATGGTGAGCAAGGGCGAG |
| eYFP_2-reverse | TTAGGATCCTTACACCTCGTTCTCG |
| NLuc-CoChR_2-forward | AATTCCCCGGGGATCCACGCGTAAGCTTTCCTTT |
| NLuc-CoChR_2-reverse | ATGGTGGCTCTAGAGAGAGATCTGCGCAAAA |

**Table 1.** List of PCR primers for plasmid construction.

| mRNA | Forward | Reserve |
|---|---|---|
| *CoChR* | CCACCAGCACATCATCATCTA | CATGGTCTCCACTTCCATCTC |
| *NLuc* | ATTGTCCTGAGCGGTGAAA | CACAGGGTACACCACCTTAAA |
| *Gapdh* | CATGGCCTTCCGTGTTCCTA | CCTGCTTCACCACCTTCTTGAT |
| *Rpl32* | CTGGAGGTGCTGCTGATGT | GGGATTGGTGACTCTGATGG |

**Table 2.** List of qRT-PCR primers.

### List of Online Supplemental Figures

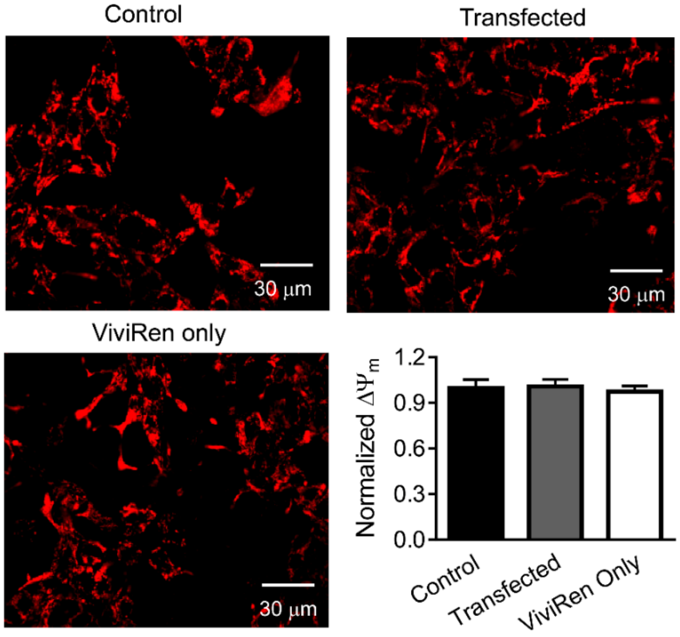


**Figure S1.** Neither mLumiOpto plasmid transfection nor ViviRen induction alone had a significant effect on cancer cell mitochondrial membrane potential ($\Delta\Psi_m$). n=4/group.

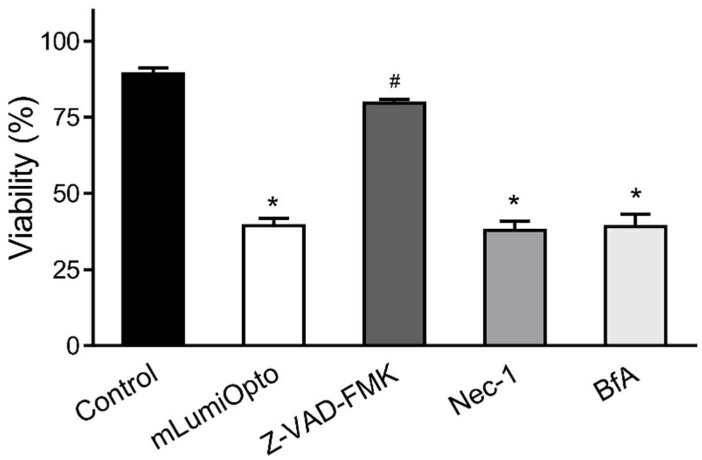


**Figure S2.** Z-VAD-FMK (a pan-caspase inhibitor) significantly alleviated mLumiOpto-mediated cell death in GBM U251 cells, whereas Nec-1 (a necrosis inhibitor) and BfA (an autophagy inhibitor) had no evident effect. *: P<0.05 vs. Control. #: P<0.05 vs. mLumiOpto. n=4/group.

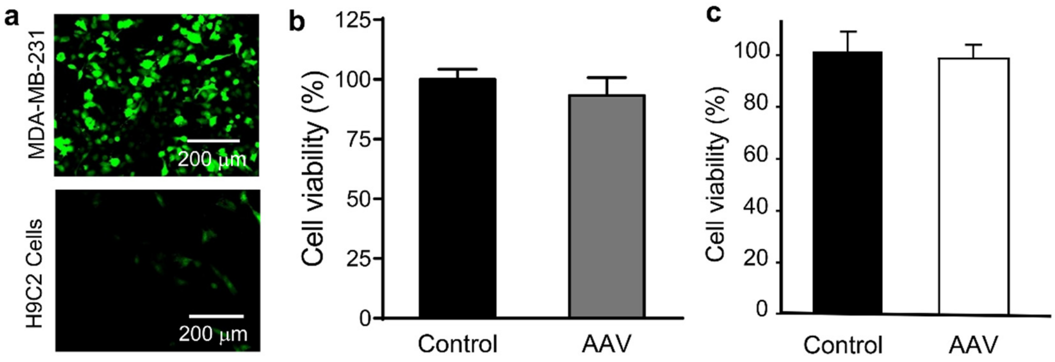


**Figure S3.** (**a**) The *cfos* promoter led to remarkably higher (>50-fold) GFP expression in cancer cells (TNBC MDA-MB-231) compared to non-cancerous cells (H9C2). (**b-c**) ViviRen had no significant effect on the viability of mLumOpto AAV-transduced non-cancer cells (HTori-3, 184B5). n=3-4/group.

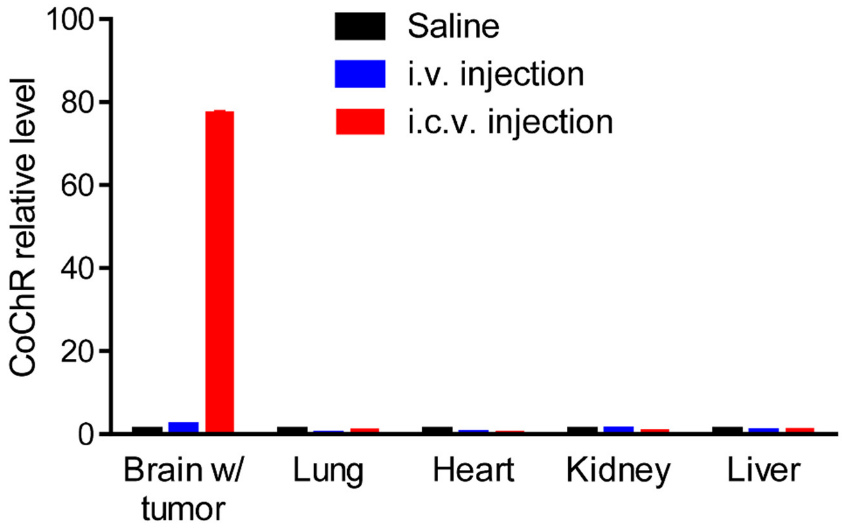


**Figure S4.** Intracerebroventricular (i.c.v.) AAV injection led to remarkably higher CoChR expression in GBM tumors compared to intravenous (i.v.) injection. n=4/group.

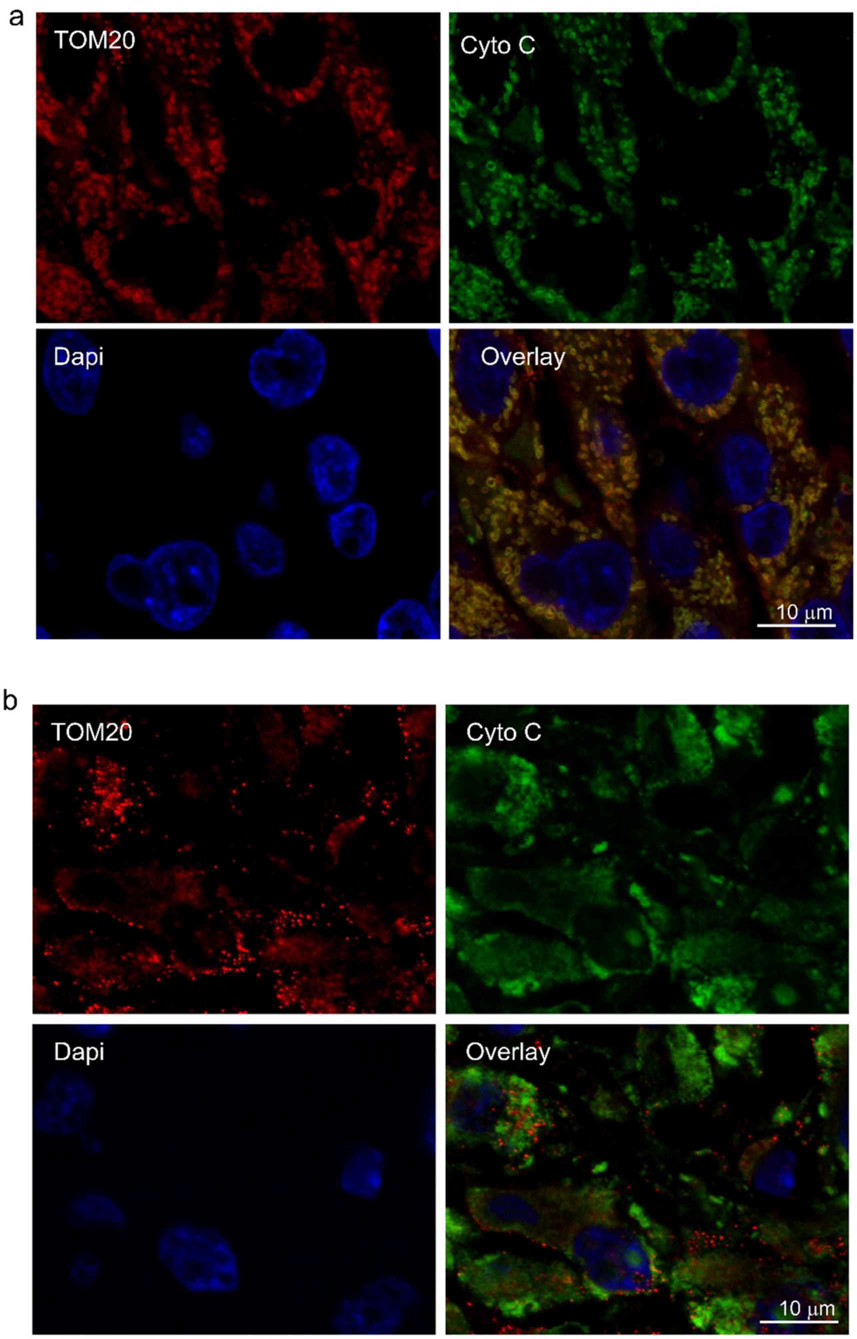


**Figure S5.** Immunofluorescence assay revealed that TOM20 overlaps with cytochrome C in the control GBM tissue (a). Cytochrome C staining becomes diffusive and TOM20 staining is fragmented in the mLumiOpto-treated GBM, indicating cytochrome C release and mitochondrial depolarization (b). Four samples were repeated in each group.

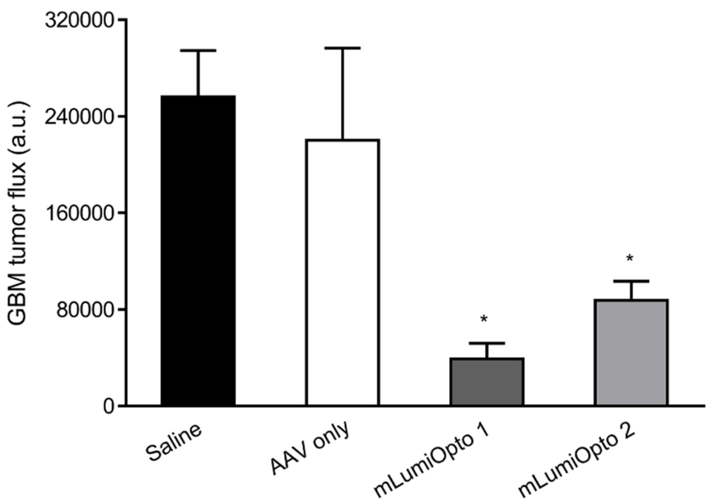


**Figure S6.** Quantification of GBM tumor bioluminescence flux collected in IVIS imaging. n=5/group.

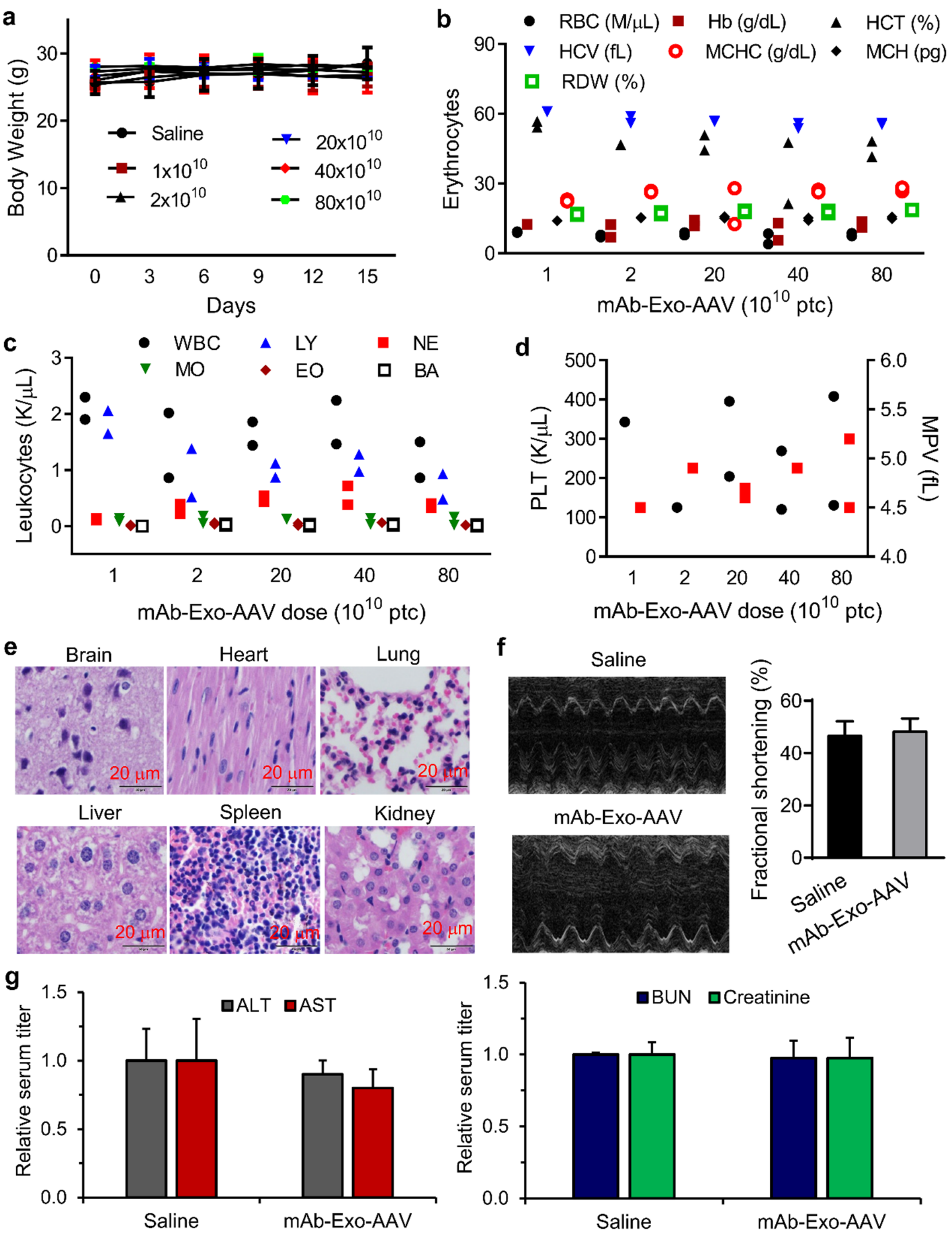


**Figure S7. Evaluation of dose tolerance and toxicity of mAb-Exo-AAV-delivered mLumiOpto.** (**a**) mLumiOpto (mAb-Exo-AAV+ViviRen) had no deleterious effect on the body weight of healthy C57BL/6J mice. (**b-d**) Whole blood analysis did not detect significant differences in erythrocytes (b), leukocytes (c) and thrombocytes (d) among various dosages. (**e**) H&E staining showed no tissue injury in the organs of high dose mLumiOpto-treated mice. (**f**) Echocardiography showed normal cardiac contractile function in healthy mice exposed to mLumiOpto. (**g**) Serum titration revealed no obvious difference in serum ALT and AST (liver markers) and BUN and creatinine (kidney markers). n=3-6/group.

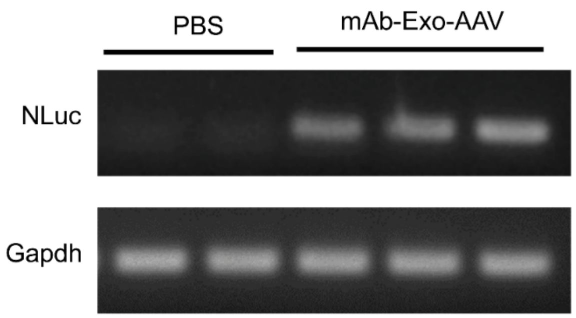


**Figure S8.** Semi-quantitative PCR revealed NLuc gene expression in the tumor of mAb-Exo-AAV mice, but not in the tumor of control (PBS) mice.

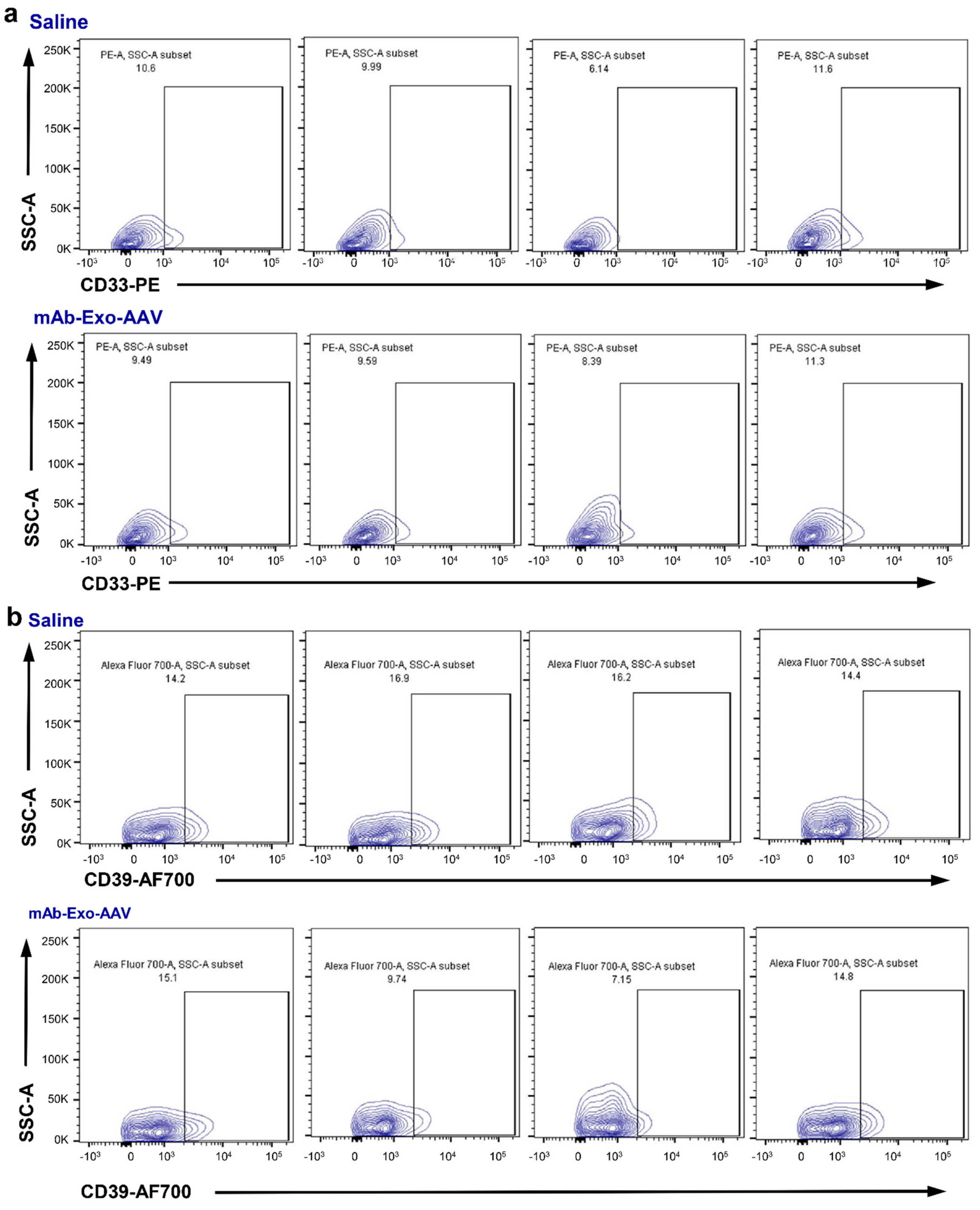


**Figure S9.** Slight or minimal regulation of immune suppression, i.e. myeloid-derived suppressor cells (a) and Treg (b), was detected in flow cytometry of fresh tumor tissues post-treatment. n=4/group.
